# Single-molecule bouncing at a liquid interface governs presynaptic enrichment

**DOI:** 10.64898/2026.11.01.763503

**Authors:** Kye Kudo, Shanley F. Longfield, Alex J. McCann, Nils Schoovaerts, Julissa Sanchez-Velasquez, Leonardo A. Parra-Rivas, Subhojit Roy, Patrik Verstreken, Elizabeth Hinde, S. Sean Millard, Adekunle T. Bademosi, Frédéric A. Meunier

## Abstract

Many cellular processes rely on accumulating highly diffusive cytoplasmic proteins to a specific location within the cell. This spatial enrichment is generally thought to rely on locally immobilizing proteins or slowing diffusion through binding and molecular crowding. Paradoxically, cytoplasmic proteins that are enriched within nerve terminals remain highly diffusive^1^, raising questions about the mechanisms that govern presynaptic enrichment. Here, using *in vivo* single-molecule microscopy, we tracked the diffusion of a model cytoplasmic protein, α-Synuclein (α-Syn), and synaptic vesicles (SVs), in motor neuron presynapses of live *Drosophila* larvae. We demonstrate that α-Syn undergoes rapid and frequent ‘bouncing’ events at a presynaptic interface, locally impeding escape from the presynapse. We measured escape probabilities at the interface, and show that impeding escape, without dramatically slowing diffusion, is sufficient to govern presynaptic enrichment. The interface is liquid-like and coincides with the boundary of the SV cluster, which is a necessary scaffold for the presynaptic enrichment of α-Syn. Our results provide direct evidence of interfacial reflectivity at the interface of liquid-like compartments, and that this phenomenon can govern the subcellular enrichment of proteins and small organelles at the level of individual particles.

## Main

The presynapse is enriched with rapidly diffusing cytoplasmic proteins that are essential for neurotransmitter release, despite being freely continuous with the axonal cytoplasm^2^. While concentrations^2^ and average diffusion coefficients^1^ of cytoplasmic proteins in the synapse have been estimated, the single-molecule dynamics of presynaptic proteins that govern their enrichment within synapses remains an open question.

Here, we address this question by tracking individual proteins within the presynapse. As a model, we chose α-Syn, a cytoplasmic protein that is presynaptically enriched^2,3^ to regulate SV recycling^4–6^. To label α-Syn for single-particle tracking with photoactivated localization microscopy (sptPALM), we transfected α-Syn knock-out (KO) primary mouse hippocampal neurons with α-Syn-mEos2 (**Fig. 1a**). We identified presynapses by co-transfecting neurons with VAMP2-pHluorin, which allows for single-SV imaging via sparse labelling with anti-GFP Atto647-nanobodies^7^. Dual-colour single-molecule microscopy and tracking in live neurons (DIV 21; **Fig. 1b**) revealed a high local density of α-Syn-mEos2 molecules at presynapses, where α-Syn-mEos2 remains mobile (**Fig. 1b**).

**Figure 1.**
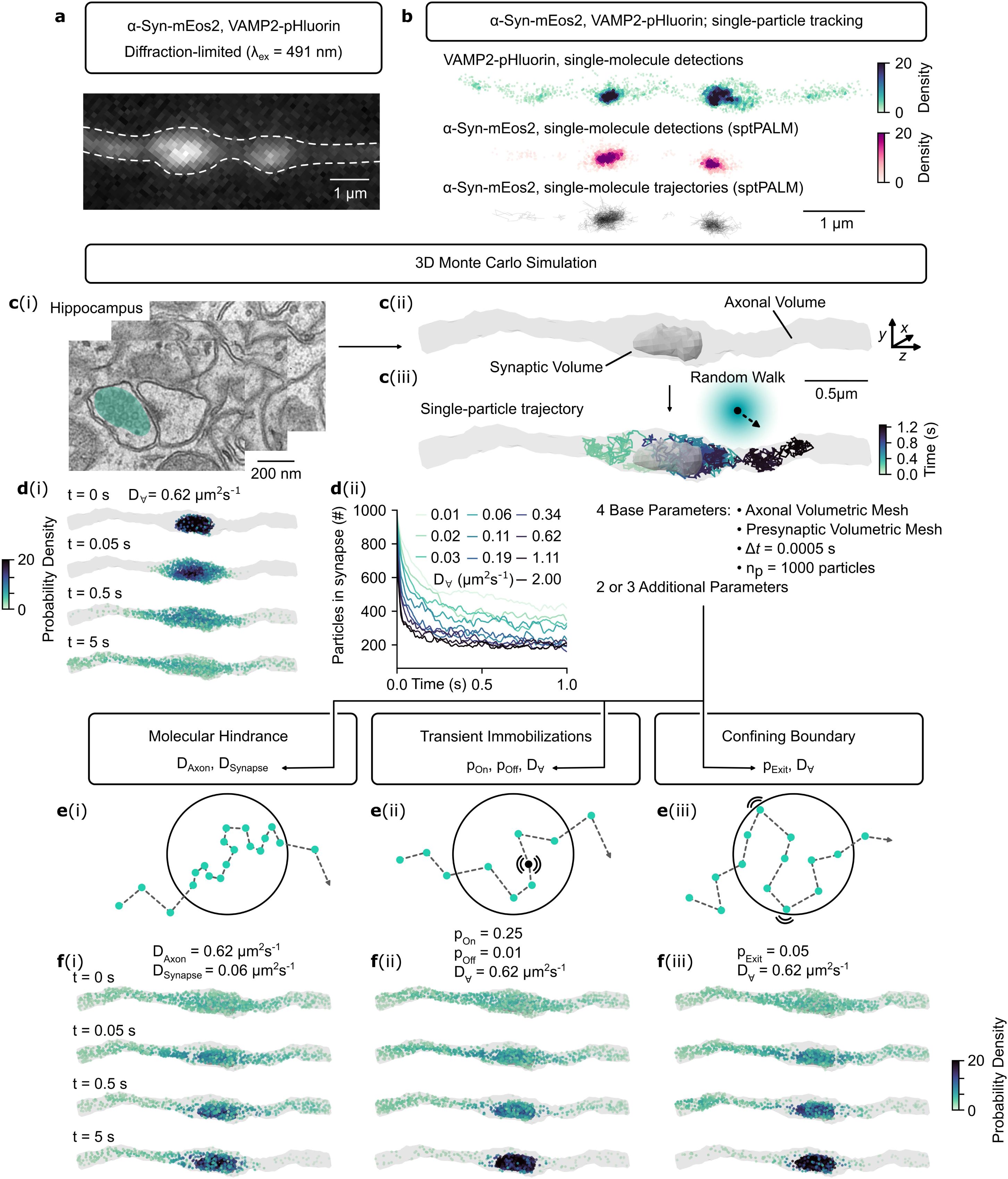
Possible models for cytoplasmic protein enrichment at the presynapse. (**a**) Representative diffraction-limited TIRF image of an axon from an α-Syn KO primary mouse hippocampal neuron co-expressing α-Syn-mEos2 and VAMP2-pHluorin. (**b**) Single-molecule localization map of (top; sdTIM) VAMP2-pHluorin and (middle; sptPALM) α-Syn-mEos2 (coloured by kernel density estimate), and (bottom) single-molecule trajectories of α-Syn-mEos2, from the axonal segment shown in **a**. (**c**) Monte Carlo simulations of single-particle cytoplasmic diffusion (**Methods**). (**c**i, ii) Volume of an axon with a presynapse generated from ssTEM acquisitions of the mouse CA1 hippocampal area. The synaptic volume (**c**i, aqua) was specified as the volume along the axon which contained SVs. (**c**iii) Representative single-particle trajectory, where diffusive motion was simulated via a random-walk. (**d**) Simple diffusion of 1000 particles starting within the presynaptic volume; (**d**i) evolution of simulation where D_∀_ = 0.62 µm^2^/s (D_∀_; global diffusion coefficient for all particles), each point represents the location of a single particle; (**d**ii) number of particles in the synapse over time (rolling average window of 30 timesteps). (**e**) Schematic illustrating possible models of presynaptic enrichment. (**e**i) Molecular hindrance: particles diffusing in the synaptic volume have a reduced diffusion coefficient compared to the rest of the axon (D_Synapse_ ≤ D_Axon_). (**e**ii) Transient immobilizations: particles in the synaptic volume have a probability (p_On_) of becoming immobilized and a probability (p_Off_) of becoming released from immobilization, with a global diffusion coefficient (D_∀_). (**e**iii) Confining boundary model: particles in the synaptic volume have the probability (p_Exit_) of escaping the synaptic volume if an exiting step is made, with a global diffusion coefficient (D_∀_). (**f**) Evolution of the simulation for each model (i), (ii) and (iii) over time, with 1000 particles initially distributed randomly along the axon. The input parameters are as displayed; each point represents the location of one particle.

### Possible mechanisms of protein diffusion and enrichment at the synapse

On short timescales, random diffusion dominates the movement of cytoplasmic proteins^1,8^. We reasoned that presynaptic enrichment could arise from three alternative mechanisms, each defined by how individual proteins move in the presynapse:

i. Molecular hindrance^9,10^: in this model, diffusing proteins are slowed at the presynapse by local crowding that sterically obstructs protein movement.
ii. Transient immobilizations^11–13^: in this model, diffusing proteins are transiently immobilized at the presynapse by interactions with a binding partner.
iii. Confining boundary^11,14^: in this model, diffusing proteins are retained in the presynapse by a ‘barrier effect’.

To assess whether these models could theoretically drive the enrichment of cytoplasmic proteins at the presynapse, we devised a Monte Carlo simulation (**Fig. 1c**). We simulated diffusion by propagating particles with independent random walks in an axon containing a presynapse reconstructed from serial-section transmission electron microscopy (ssTEM)^15^, as similarly done before^1^ (**Fig. 1c**, **Supplementary Note 1**). We propagated particles at mobilities spanning 0.01 - 2 µm^2^/s, capturing a broad range of estimated diffusion coefficients for cytoplasmic proteins along the axon and presynapse^1,16^. Under simple diffusion, particles rapidly diffuse out of the presynapse within a second (**Fig. 1d**), indicating that a mechanism of presynaptic enrichment is needed.

We found that the three models (**Fig. 1e**), can drive enrichment whilst maintaining internal mobility and exchange (i.e., flux), depending on the input parameters (**Fig. 1f** and **Extended Data Fig. 1**). While these models are not mutually exclusive, we suspected that there is a predominant mechanism that governs presynaptic diffusion and enrichment. To investigate, we leveraged presynaptic boutons of the *Drosophila* larval neuromuscular junction (NMJ), which are considerably larger (2 - 5 µm) than the presynapses of cultured neurons, allowing us to resolve single-molecule movements within local regions of the presynapse.

### α-Syn is enriched at motor neuron presynapses

Muscle-13 of *Drosophila* larvae is innervated by type-Ib boutons of the glutamatergic motor neuron, MN13-Ib^17^, which we used for this study. We expressed α-Syn-mEos3.2 in motor neurons using the GAL4/UAS system^18^ and imaged MN13-Ib boutons in live larvae by confocal microscopy; MN13-Ib boutons are readily identified as large swellings along the axon terminal^17^. *Drosophila* lack an α-Syn homologue and therefore provide a null background to study its organization. We found that α-Syn-mEos3.2 localized to discrete peripheral clusters in MN13-Ib boutons (**Fig. 2a**). We quantified this peripheral clustering by extracting intensity profiles across boutons and calculating the ratio of peripheral-versus-central intensity (**Fig. 2a-c**). α-Syn-mEos3.2 showed significant peripheral enrichment (ratio > 1) in 66 % of boutons (ratio x̅ = 1.6). In comparison, when the mEos3.2-tag is expressed on its own, it was uniformly distributed within boutons (ratio x̅ = 1.0), indicating that the boutons’ cytoplasm is broadly accessible to freely diffusive molecules^19^ (**Fig. 2a-c**). Together, α-Syn-mEos3.2 is selectively enriched at peripheral clusters within MN13-Ib boutons.

**Figure 2.**
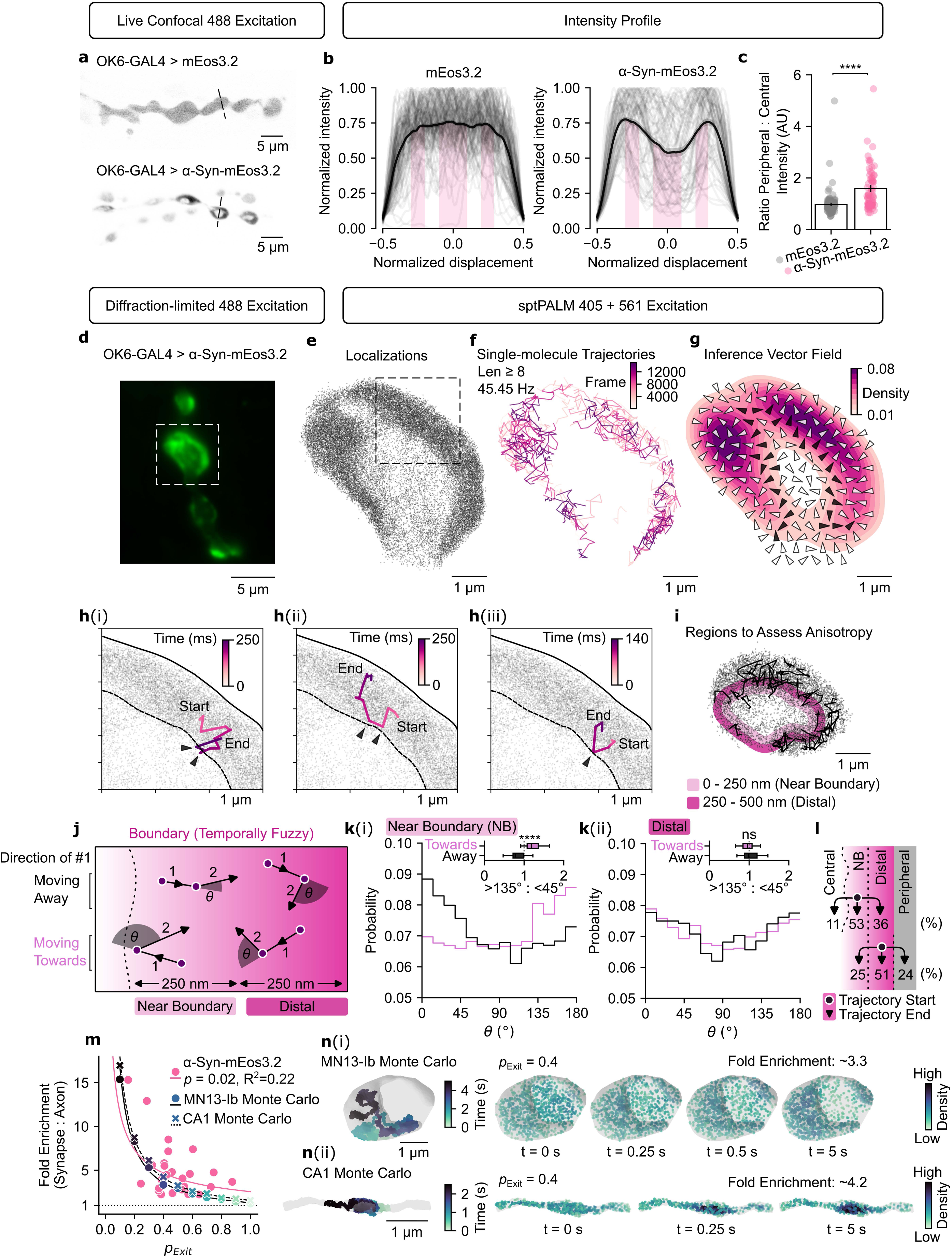
α-Syn molecules ‘bounce’ at a boundary in the presynapse. (**a**) Representative confocal images of MN13-Ib from live larvae expressing mEos3.2 (top) or α-Syn-mEos3.2 (bottom). (**b**) Fluorescence intensity profiles across boutons expressing mEos3.2 (left) or α-Syn-mEos3.2 (right); individual boutons (grey) and mean (black). (**c**) Ratio of peripheral to central intensity acquired from intensity profiles in **b** (mEos3.2, *n* = 118 boutons from 4 larvae; α-Syn-mEos3.2, *n* = 73 boutons from 3 larvae; two-sided Mann-Whitney U test). (**d**) Oblique illumination of α-Syn-mEos3.2 at representative MN13-Ib boutons. (**e**, **f**, **g**) SptPALM of α-Syn-mEos3.2 in a representative MN13-Ib bouton indicated in **d** acquired over 330 s at 45.45 Hz showing (**e**) single-molecule localizations, (**f**) single-molecule trajectories, and (**g**) vector field of local directional bias computed from single-molecule trajectories overlayed on top of the kernel density estimate of single-molecule detections. Vectors near the boundary between the central axoplasm and peripheral cluster are coloured black. (**h**) Representative single-molecule trajectories of α-Syn-mEos3.2 display hindered motion at the boundary; arrows indicate locations in the trajectory where α-Syn-mEos3.2 appears to ‘bounce’. (**i**) For quantification of anisotropy, a region at the boundary is specified and dilated outwards by 250 nm (*near boundary*), and an additional 250 nm (*distal*). (**j**) Graphical representation of trajectories near and distal to the boundary. If the boundary impedes diffusion, a molecule near the boundary that takes a step towards the boundary should be more likely to ‘bounce back’ (*θ* >135°) in the following step, whilst molecules with unhindered motion should show no preference. (**k**) Measurements of angles made within trajectories that are (**k**i) near the boundary and (**k**ii) distal to the boundary, grouped by molecules that move towards or away from the boundary in an initial step. Inset boxplots are the ratio of *θ* that are >135° (backward step) versus <45° (forward step), calculated per bouton (*n* = 25 boutons from 7 larvae, 8284/12010/14282/18732 steps for near-away/near-towards/distal-away/distal-towards; two-sided paired Wilcoxon signed-rank test). (**l**) Percentages of molecules that started within the near-boundary (NB) and distal regions that ended more central or peripheral. (**m**) Fold enrichment as a function of *p_Exit_*, measured experimentally for α-Syn-mEos3.2 (pink), and derived using Monte Carlo simulations (*D_Axon_* = 0.41 µm^2^/s, *D_Synapse_* = 0.31 µm^2^/s, coloured by *p_Exit_*), lines correspond to respective fits. (**n**) Monte Carlo simulations (left) trajectory and (right) particle density, at *p_Exit_* = 0.4, for (**n**i) MN13-Ib and (**n**ii) CA1 boutons. Data shown as mean ± SEM; *p*-values indicated as: ****, *p* ≤ 0.0001; ns, *p* > 0.05.

### α-Syn remains mobile within the presynaptic compartment

We next used *in vivo* sptPALM to track individual α-Syn-mEos3.2 molecules in MN13-Ib boutons of live larvae^20,21^ (**Supplementary Movie 1**). Super-resolved detection maps confirmed that α-Syn-mEos3.2 was organised into peripheral clusters, with a 5.03 ± 0.65-fold enrichment (median, ∼4.1) in single-molecule localization density in the peripheral cluster (1495 ± 168 detections/µm^2^) compared to central axoplasm (345 ± 44 detections/µm^2^; **Fig. 2de**, **Extended Data Fig. 2a**). α-Syn-mEos3.2 remained mobile within peripheral clusters (apparent diffusion coefficient, *D* = 0.31 ± 0.04 μm^2^/s), although mobility was slightly higher in the central axoplasm (0.41 ± 0.05 μm^2^/s; **Fig. 2f, Extended Data Fig. 2b-e**). We validated our estimated diffusion coefficients with line-scanning fluorescence correlation spectroscopy (FCS)^22^ (**Extended Data Fig. 3a-e**). FCS measurements were modestly higher overall but remained consistent with the relative increase of mobility in the central axoplasm (0.94 ± 0.11 μm^2^/s) compared to the peripheral cluster (0.62 ± 0.07 μm^2^/s). The presynaptic diffusion coefficient of the mEos3.2-tag, expressed on its own, was 13.81 ± 1.06 μm^2^/s (**Extended Data Fig. 3e**). Since the molecular weights of α-Syn-mEos3.2 and mEos3.2 are of the same order of magnitude, the significantly lower mobility of α-Syn-mEos3.2 is likely to be independent of molecular crowding^16^, and may instead reflect interactions of α-Syn. Next, we estimated the fraction of immobile molecules^23^, which often reflects long-lived binding interactions. ∼23% and ∼22% of α-Syn-mEos3.2 molecules were immobile in the central axoplasm and peripheral cluster, respectively, suggesting that long-lived binding events occur to the same extent in both intra-synaptic regions (**Extended Data Fig. 2f**). It is possible that ‘immobilizing’ events of α-Syn are ultra-transient and beyond our fast temporal resolution, in this case, ensembles of such events are captured within the apparent diffusion coefficients^24^. Given this, we asked whether the difference in diffusion coefficients between the intra-synaptic regions could account for peripheral enrichment using our Monte Carlo simulation. We found that the reduced mobility in the peripheral cluster, is unable to account for the 5-fold enrichment in the periphery (**Extended Data Fig. 3f,g**). Together, these results suggest that neither molecular crowding nor transient immobilizations are predominant mechanisms for enriching α-Syn-mEos3.2 at the presynapse.

### A ‘barrier effect’ governs the presynaptic enrichment of α-Syn

We next examined whether a ‘barrier effect’ may explain peripheral enrichment of α-Syn-mEos3.2. First, we used single-molecule trajectories to determine whether molecules show local directional biases^25,26^. This revealed that α-Syn-mEos3.2 molecules near the axoplasm-cluster boundary experienced a directional bias that was directed away from the central axoplasm (**Fig. 2g, Extended Data Fig. 2h**). α-Syn-mEos3.2 molecules within the peripheral cluster that approached the axoplasm-cluster boundary frequently appeared to ‘bounce back’, which could account for this directional bias (**Fig. 2h**, **Extended Data Fig 2j**). To investigate further, we assessed for single-molecule ‘bouncing’ events by measuring the angles (*θ*) made within trajectories near the axoplasm-cluster boundary, which we specified as a 250 nm-wide region (**Fig. 2i,j**). We found that when a molecule near the boundary takes a step from the cluster *towards* the central axoplasm, it was more likely to ‘bounce back’ (*θ* > 135°) in its following step, this is illustrated in the *θ* histogram as well as the ratio between forward (*θ* < 45°) and backward angles (*θ* > 135°) (**Fig. 2k**i; *near boundary*). This anisotropic movement was not observed further away (*distal*) from the boundary (**Fig. 2k**ii). This indicates that a ‘barrier effect’ is experienced by individual molecules. In addition to molecules that ‘bounce’, α-Syn-mEos3.2 trajectories that enter and exit the peripheral cluster are also observed (**Extended Data Fig 2k,l**), however we expected that if ‘bouncing’ events occurred frequently and impeded diffusion, only a small proportion of molecules would escape. Of those single-molecule trajectories that began within the near-boundary region, ∼11% exited the cluster to a more central position, whilst ∼36% of trajectories remained within the cluster in a more peripheral position (**Fig. 2l**). In comparison, molecules whose trajectories started in the distal region had equal probabilities of ending more central (∼25%) or more peripheral (∼24%) (**Fig. 2l**). This indicates that a ‘barrier effect’ impedes the diffusion of molecules out from the peripheral cluster. To determine if this ‘barrier effect’ could account for peripheral enrichment, we estimated the probability, *p_Exit_*, of a molecule near the boundary escaping from the peripheral cluster (on ∼10-100 millisecond timescales; *p_Exit_* = 1, free escape; *p_Exit_* = 0, no escape) based on the above percentages, obtaining *p_Exit_* = 0.48 ± 0.03 (median, ∼0.43; **Methods**). We plotted the fold enrichment in the peripheral cluster against estimated *p_Exit_* for each bouton (*n* = 25 boutons) and found that peripheral enrichment inversely correlates with *p_Exit_* (power function, a × *p_Exit_^b^*, **Fig. 2m**; *pink*). This correlation was similarly observed in Monte Carlo simulations (model *iii*) when simulating across different values of *p_Exit_*, within a MN13-Ib bouton and CA1 axon-segment (**Fig. 2m,n**). Together, these results indicate that a ‘barrier effect’ governs the presynaptic enrichment of α-Syn-mEos3.2. Interestingly, the correlation obtained from the simulations in MN13-Ib and CA1 boutons are similar, suggesting that a ‘barrier effect’ may allow for universal tuning of synaptic enrichment in a way that is independent of synapse morphology (**Fig. 2m**).

### SVs experience a ‘barrier effect’ at the presynapse

SVs are clustered at the periphery of *Drosophila* NMJ boutons^27,28^. This can be observed in images of motor neuron boutons acquired on an electron microscope (**Fig. 3a**). Synaptotagmin 1 (Syt1), an abundant transmembrane protein of SVs^29^, colocalized with α-Syn-mEos3.2 in MN13-Ib boutons (**Extended Data Fig. 4**). We therefore suspected that SVs contribute to the presynaptic enrichment of α-Syn, and set out to characterize the organization and dynamics of SVs.

**Figure 3.**
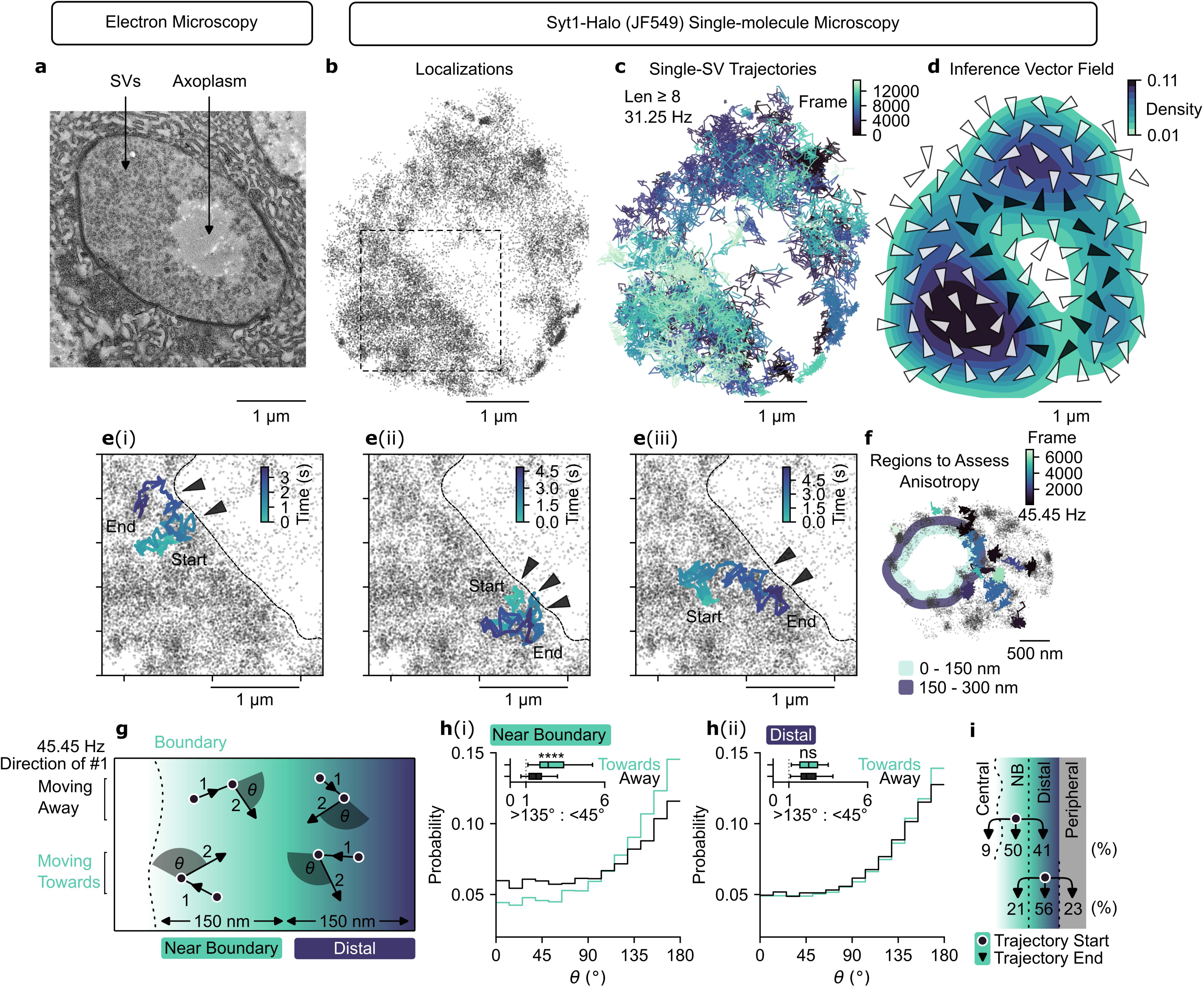
Synaptic vesicles ‘bounce’ at a boundary in the presynapse. (**a**) Representative image of a *Drosophila* motor neuron bouton captured on an electron microscope. (**b**, **c**, **d**) Single-molecule microscopy of Syt1-Halo (JF549) in a representative MN13-Ib bouton, this acquisition was made over 480 s at 31.25 Hz. (**b**) Single-molecule SV localizations, (**c**) trajectories, and (**d**) local directional biases computed from single-molecule trajectories, overlayed on top of the kernel density estimate of localizations; vectors near the boundary of the central axoplasm and the peripheral SV cluster are coloured black. (**e**) Representative single-molecule trajectories of SVs displaying hindered motion at the boundary between the central axoplasm and the peripheral SV cluster from the region shown in **b**. Black arrows indicate the location in the trajectory where SV motion appears hindered across the boundary (for more representative acquisitions, see **Extended Data** Fig. 5). (**f**) Representative example of regions of interest used for anisotropic quantification; *near-boundary* (0 - 150 nm; light blue) and *distal* (150 - 300 nm; dark blue). (**g**) Graphical representation of trajectories near and distal to the boundary. If the boundary alters diffusion, an SV near the boundary that takes a step towards the boundary should be more likely to ‘bounce back’ (θ >135°) in the following step. (**h**) Measurements of angles made within trajectories that are (**h**i) near the boundary and (**h**ii) distal to the boundary for molecules that move towards or away from the boundary in their first step. Inset boxplots are the ratio of steps that are >135° (backward step) versus <45° (forward step), calculated per bouton (*n* = 38 boutons from 9 larvae, 18616/29048/39825/51059 steps for near-away/near-towards/distal-away/distal-towards; two-sided paired Wilcoxon signed-rank test). (**i**) Percentages of trajectories that started within the near-boundary (NB) and distal regions that ended more central or peripheral. Data shown as mean ± SEM; *p*-values indicated as: ****, *p* ≤ 0.0001; ns, *p* > 0.05.

We expressed Syt1-EGFP in MN13-Ib boutons^28^. In live larvae, Syt1-EGFP localized to the bouton periphery, whilst the EGFP-tag alone distributed uniformly (**Extended Data Fig. 5a-c**). 73% of MN-13Ib boutons exhibited peripheral enrichment (ratio > 1) of Syt1-EGFP (ratio x̅ = 1.8; EGFP, ratio x̅ = 1.0). Fluorescence Recovery After Photobleaching (FRAP) experiments of a small region in the peripheral SV cluster showed recovery of Syt1-EGFP for many (but not all) boutons confirming that SVs are largely mobile within the presynapse^28,30^ (**Supplementary Movie 2**).

To directly assess single-SV movements, we expressed Syt1-Halo in MN13-lb/MN12-Ib boutons and performed single-molecule microscopy using sparse labelling with the membrane-permeable JF549 Halo-tag ligand. Since SVs are ∼40 nm in diameter, our acquisitions capture the translational motion of the SVs whilst the circumferential movements of Syt1-Halo on the SV membrane are occluded by localization precision^7,16^. Single-particle localizations of Syt1-Halo (JF549) at MN13-Ib/MN12-Ib boutons were detected at the periphery in 54 of 75 boutons captured, confirming the peripheral clustering of SVs (**Fig. 3b** and **Extended Data Fig. 5g**). The diffusion coefficient of SVs were lower in boutons that displayed peripheral clustering (0.053 ± 0.006 µm^2^/s; consistent with previous measurements^7,16,31^) compared to those without (0.121 ± 0.015 µm^2^/s; **Extended Data Fig. 5g-k**). We note that whilst diffusion coefficients were dramatically lower than α-Syn-mEos3.2, some SVs remained mobile in the peripheral cluster (0.052 ± 0.007 µm^2^/s; immobile fraction ∼59%) (**Fig. 3c** and **Extended Data Fig. 5k**). We suspected that SV movement may be altered at the cluster’s interface and sought to assess whether SVs experience a ‘barrier effect’. Maps of local directional biases revealed that SVs near the cluster’s boundary, experienced a directional bias that was directed away from the central axoplasm (**Fig. 3d** and **Extended Data Fig. 5l-m**). Individual SVs frequently appeared to ‘bounce back’ as they approached the central axoplasm (**Fig. 3e** and **Extended Data Fig. 5o**). As before, we quantitively assessed for ‘bouncing’ events at a near-boundary region (**Fig. 3f,g**). Angles (*θ*) made within SV trajectories were often > 135° in the peripheral cluster (*distal*), which can be explained by the large immobile fraction and diffusion rates on the order of localization precision (**Fig. 3h**ii). However, we found that SVs near the boundary that take a step towards the central axoplasm, had an increased probability to ‘bounce back’ (θ > 135°) (**Fig. 3h**i). This indicates that a ‘barrier effect’ is experienced by individual SVs. Of those single-particle trajectories that began within the near-boundary region, ∼9% escaped the cluster and ended in the central axoplasm, whilst ∼91% of trajectories remained in the peripheral cluster (**Fig. 3i**). Together, these results suggest that, like α-Syn, a ‘barrier effect’ at the cluster’s boundary impedes the diffusion of SVs out from the peripheral cluster.

### ‘Barrier effect’ is a result of a presynaptic liquid-like interface

We asked what interactions are responsible for this ‘barrier effect’. In cellular structures formed via phase separation, proteins establish multivalent interactions^32^. To escape, a protein must break multiple interactions, which is energetically unfavourable. In effect, it has been predicted that the interface of a phase-separated compartment behaves as an energetic barrier for diffusion, favouring internal retention^11,33,34^. Our ‘bouncing’ observations are consistent with this behaviour, and confocal timelapse imaging revealed wave-like fluctuations of the peripheral cluster’s interface^35,36^ (**Extended Data Fig. 3m-o** and **Extended Data Fig. 5d-f** and **Supplementary Movie 3**), moreover α-Syn has been shown to phase-separate *in vitro*^37^. We thus suspected the involvement of phase-separation. 1,6-hexanediol (1,6-HD) can dissolve phase-separated structures that are stabilized by hydrophobic interactions^38,39^. 1,6-HD (4%) abolished the peripheral enrichment of α-Syn-mEos3.2, which was reversed upon washout (**Fig. 4a-c**). 1,6-HD also disrupted the diffusional bias and increased the mobility of α-Syn-mEos3.2 (**Fig. 4 d-f**). We also observed an increase in α-Syn-mEos2 mobility in presynapses of α-Syn KO hippocampal neurons following incubation with 1,6-HD (1%) (**Extended Data Fig. 6a-c**). The hydrophobic ‘non-amyloid-β component’ (NAC) domain is critical for α-Syn’s ability to phase separate^37^. β-Syn, the homologue of α-Syn, principally lacks residues 73-83, that constitutes part of the NACore^40^ (residues 68-78) and does not spontaneously phase separate^37,41,42^. NACore deletion (α-Syn^ΔNAC^-mEos3.2) dramatically impaired peripheral clustering and increased presynaptic mobility of α-Syn in MN13-Ib boutons (**Extended Data Fig. 6d-h**). These results suggest that the presynaptic enrichment of α-Syn-mEos3.2 may depend on interactions typically associated with liquid-like phase-separated compartments.

**Figure 4.**
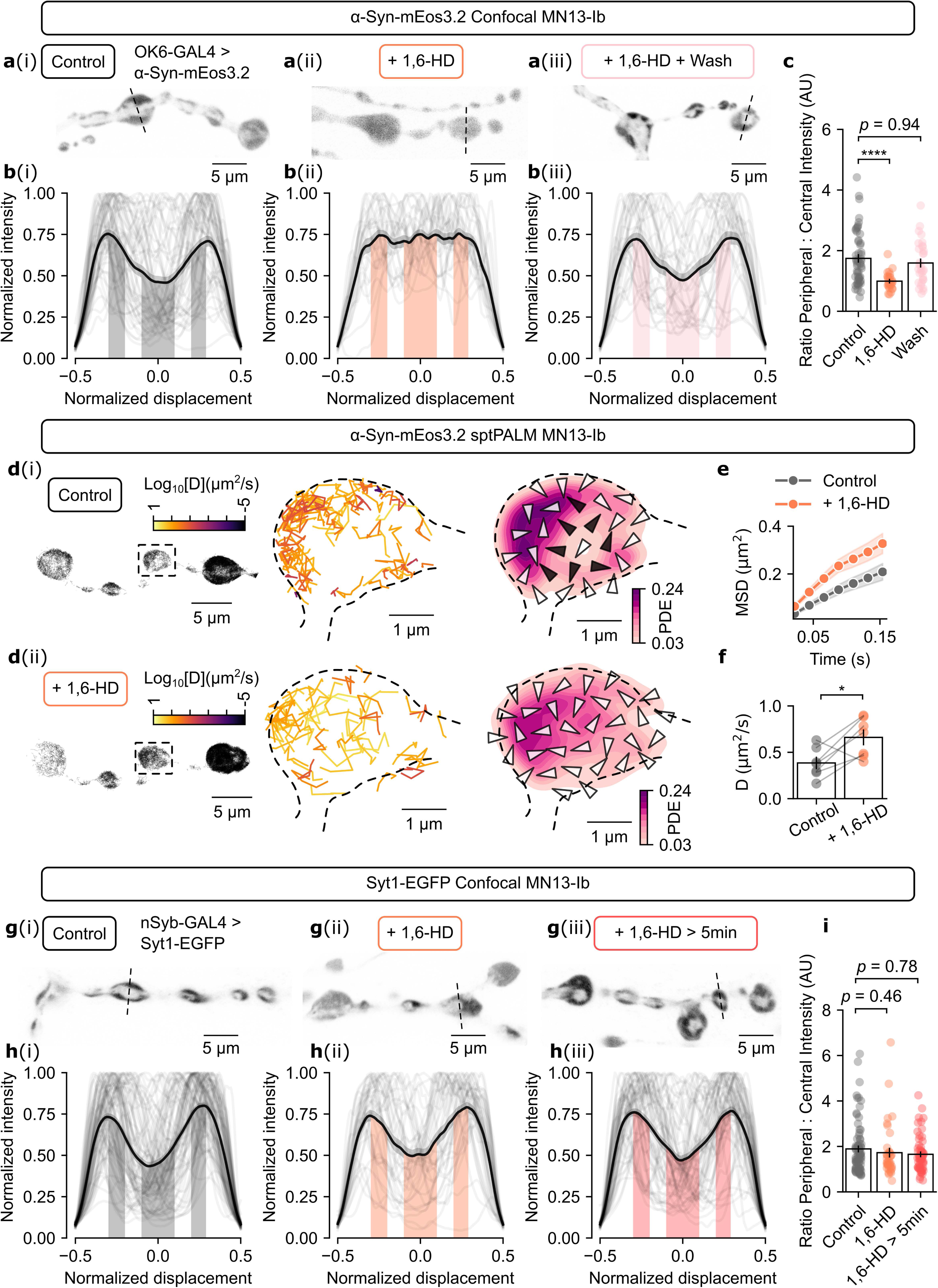
1,6-HD disrupts the clustering of α-Syn, but not of SVs. (**a**) Representative confocal images of MN13-Ib from live larvae expressing α-Syn-mEos3.2, under (i) vehicle (modified HL3) control, (ii) addition of 4% 1,6-HD for < 5 min, and (iii) addition of 1,6 HD (4%; 5 min) followed by modified HL3 wash-off (≥10 min). (**b**) Fluorescence intensity profiles of α-Syn-mEos3.2 across MN13-Ib boutons, under conditions described for **a**. Individual boutons (grey) and mean of the intensity profile (black). (**c**) Ratio of peripheral versus central intensity from intensity profiles in **b** (Control, *n* = 54 boutons from 3 larvae; 1,6-HD, *n* = 26 boutons from 3 larvae, *p =* 0.00006; Wash, *n* = 31 boutons from 3 larvae, *p* = 0.94; Dunn’s test, Dunn-Šidák correction,). (**d**) Single-molecule microscopy of α-Syn-mEos3.2 at MN13-Ib in vehicle control and 1,6-HD (4%) for < 5 min; (left) zoomed-out single-molecule trajectory map, (middle) single-molecule trajectory map from inset (coloured by diffusion coefficient), (right) vector field of local directional bias overlayed with kernel density estimate of single-molecule detections; (**d**i) for vehicle control and (**d**ii) 1,6-HD (4%). (**e**) MSD-Δ*t* plot for vehicle control and 1,6-HD (4%) for < 5 min. (**f**) Apparent diffusion coefficients calculated from MSD-Δ*t* plot (Control, *n* = 7 MN13-Ib NMJs from 7 larvae; 1,6-HD, *n* = 7 MN13-Ib NMJs from 7 larvae; two-sided paired Wilcoxon signed-rank test, *p* = 0.047). (**g**) Representative confocal images of MN13-Ib from live larvae expressing Syt1-EGFP, under (i) vehicle (modified HL3) control, (ii) addition of 4% 1,6-HD for < 5 min, and (iii) addition of 1,6 HD (4%) > 5min (and up to 11 min). (**h**) Fluorescence intensity profiles of Syt1-EGFP across MN13-Ib boutons, under conditions described for **g**. Individual boutons (grey) and mean of the intensity profile (black). (**i**) Ratio of peripheral versus central intensity from intensity profiles in **h** (Control, *n* = 83 boutons from 5 larvae; 1,6-HD, *n* = 39 boutons from 5 larvae; 1,6-HD > 5 min, *n* = 61 boutons from 6 larvae, Dunn’s test, Dunn-Šidák correction). Data shown as mean ± SEM; *p*-values indicated as: *, p ≤ 0.05; ****, *p* ≤ 0.0001; *ns*, not significant *p* > 0.05.

Interestingly, 1,6-HD (4%) did not abolish the peripheral clustering of SVs (**Fig. 4g-i**), consistent with ref.^43^, but see also ref.^44^. Synapsin contributes to clustering SVs^45^. In synapsin-null *Drosophila* motor neurons, peripheral SV clustering^46–48^ and NMJ outgrowth^49^ are disrupted. The EGFP-tagged *Drosophila* synapsin isoform (EGFP-synapsin), that has proline-rich regions and conserved ‘C’ domain^50^, can rescue the latter phenotype^49^. EGFP-synapsin is enriched at the periphery of MN13-Ib presynaptic boutons (**Extended Data Fig. 7**). 1,6-HD (4%) abolished the peripheral enrichment of EGFP-synapsin, which was reversed upon wash off (**Extended Data Fig. 7**). This suggests that in addition to synapsin, other presynaptic proteins contribute to SV clustering. To gain further insight, we performed an RNA interference (RNAi) screen against candidate genes and scored the peripheral localisation of Syt1-EGFP. 42 candidates were selected based on whether encoding proteins contained intrinsically disordered regions or were presynaptic (**Extended Data Fig. 8**). Knock-down (KD) of 11 candidates resulted in impaired clustering of SVs, including SV-associated integral membrane proteins: vesicular glutamate transporter 1 (VGlut1), Synaptogyrin, and Synaptotagmin-7. Thus, in addition to synapsin, other presynaptic proteins are required for maintaining the SV cluster, and that SVs may be actively involved as an organizer of liquid-like cytoplasmic clustering.

### SV scaffold is essential for α-Syn enrichment

We hypothesized that SVs are a scaffold that localizes the peripheral clustering of α-Syn mediating the ‘barrier effect’. If this is the case, (i) altering the interaction strength between α-Syn and SVs, and (ii) removing the SV scaffold, should disrupt the presynaptic clustering of α-Syn. The N-terminal region of α-Syn forms an α-helical coil that interacts with highly curved membrane structures, including SVs^51^. Parkinson’s disease-associated missense mutations alter the affinity of α-Syn to curved membranes *in vitro*^51^: E46K and H50Q increase affinity^52–54^, whereas A30P and G51D decrease affinity^52,53,55^. We found that α-Syn^E46K^-mEos3.2 and α-Syn^H50Q^-mEos3.2 are peripherally clustered within MN13-Ib boutons, but diffuse significantly slower than the wild-type (**Fig. 5a-c**, **Extended Data Fig. 9a,b**). α-Syn^A30P^-mEos3.2 and α-Syn^G51D^-mEos3.2 have dramatically impaired peripheral clustering and diffuse significantly faster than wild-type. Interestingly, E46K substitution, but not H50Q, caused the peripheral clustering of α-Syn to be insensitive to 1,6-HD (**Extended Data Fig. 9c-h**). Together, the binding affinity of α-Syn to curved membranes appears to correlate with the recruitment to the peripheral cluster.

**Figure 5.**
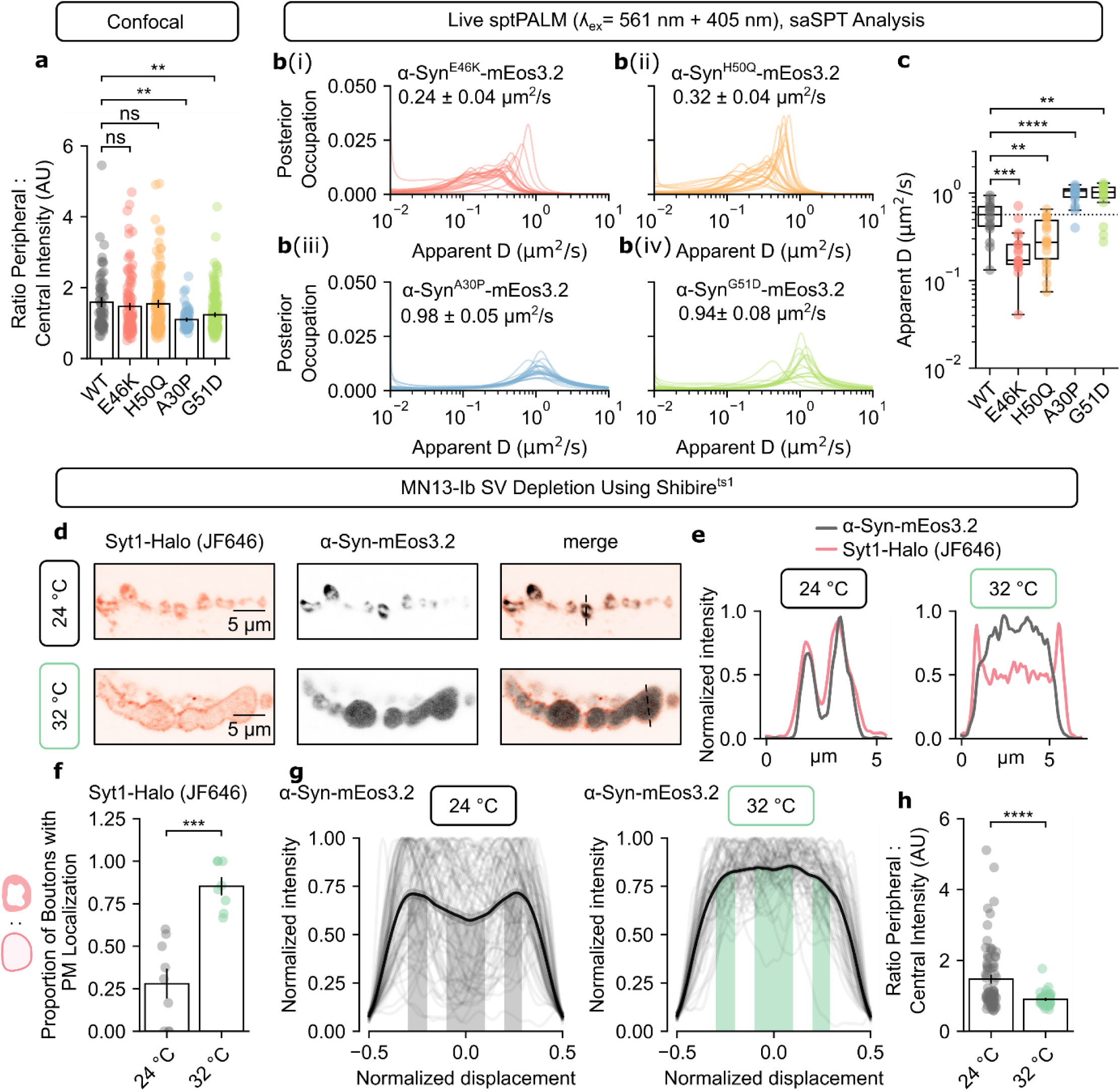
SV scaffold is essential for α-Syn enrichment. (**a**) Ratio of peripheral to central intensity acquired from intensity profiles across MN13-Ib boutons in live *Drosophila* larvae expressing different α-Syn mutants under OK6-GAL4 (**Extended Data** Fig. 9; α-Syn-mEos3.2, *n* = 73 boutons from 3 larvae; α-Syn^E46K^-mEos3.2, *n* = 110 boutons from 4 larvae; α-Syn^H50Q^-mEos3.2, *n* = 128 boutons from 5 larvae; α-Syn^A30P^-mEos3.2, *n* = 65 boutons from 3 larvae; α-Syn^G51D^-mEos3.2, *n* = 156 boutons from 6 larvae; two-sided Mann-Whitney U Test, Dunn-Šidák correction). (**b**) Distribution of diffusion coefficient inferred using saSPT from single-molecule trajectories of (i) α-Syn^E46K^-mEos3.2, (ii) α-Syn^H50Q^-mEos3.2, (iii) α-Syn^A30P^-mEos3.2, and (iv) α-Syn^G51D^-mEos3.2. (**c**) Median apparent diffusion coefficients obtained from saSPT (α-Syn-mEos3.2, *n* = 18 NMJs from 8 larvae; α-Syn^E46K^-mEos3.2, *n* = 21 NMJs from 6 larvae; α-Syn^H50Q^-mEos3.2, *n* = 23 NMJs from 8 larvae; α-Syn^A30P^-mEos3.2, *n* = 21 NMJs from 6 larvae; α-Syn^G51D^-mEos3.2, *n* = 16 NMJs from 6 larvae; two-sided Mann-Whitney U Test, Dunn-Šidák correction). (**d**) Representative confocal images of MN13-Ib NMJs from live Shibire^ts1^ larvae expressing Syt1-Halo (labelled with JF646; 200 nM) and α-Syn-mEos3.2, after high K^+^ stimulation (1 min) and recovery period at 24 °C and 32 °C. (**e**) Representative intensity profiles of Syt1-Halo (JF646) and α-Syn-mEos3.2 across boutons shown in **d**. (**f**) Proportion of boutons with significant Syt1-Halo (JF646) fluorescence at the plasma membrane (PM) within MN13-Ib NMJ chain, manually determined by confocal acquisitions of Syt1-Halo (JF646) (24 °C, *n* = 9 NMJs from 3 larvae; 32 °C, *n* = 8 NMJs from 3 larvae). (**g**) Profiles of fluorescence intensity of α-Syn-mEos3.2 across MN13-Ib boutons in Shibire^ts1^ mutants after high K^+^ stimulation (1 min) and recovery period at 24 °C and 32 °C. Intensity profiles of individual boutons (grey) and mean intensity profile (black). (**h**) Ratio of peripheral to central intensity acquired from intensity profiles (24 °C, *n* = 76 boutons from 3 larvae; 32 °C, *n* = 62 boutons from 3 larvae). Data shown as mean ± SEM; *p*-values indicated as: **, p ≤ 0.01; ***, p ≤ 0.001, ****, *p* ≤ 0.0001; *ns*, not significant *p* > 0.05.

To determine if SVs are required for the enrichment of α-Syn, we depleted SVs from MN13-Ib boutons using the temperature-sensitive dynamin mutant, Shibire^ts1^ (Shi^ts1^)^56^. In Shi^ts1^ larvae, dynamin-dependent SV endocytosis is inhibited at restrictive temperatures (> 29°C). Stimulating MN13-Ib at restrictive temperatures depletes SVs from presynapses by blocking SV recycling^57^. We imaged live Shi^ts1^ larvae expressing Syt1-Halo (JF646) and α-Syn-mEos3.2. After stimulating at restrictive temperatures, Syt1-Halo localized to the presynaptic plasma membrane, outlining the MN13-Ib boutons (**Fig. 5d,e**), indicating depletion of SVs following their fusion with the presynaptic plasma membrane (**Fig. 5f**). In these boutons, the peripheral clustering of α-Syn-mEos3.2 is abolished (**Fig. 5d-h**). At permissive temperatures (< 24°C), Syt1-Halo localized to peripheral clusters and α-Syn-mEos3.2 displayed peripheral clustering (**Fig. 5d-h**). To assess this effect in hippocampal neurons, we performed sptPALM of α-Syn-mEos2 in α-Syn KO hippocampal neurons following incubation with Dyngo-4a, a dynamin inhibitor^58–60^. The mobility of presynaptic α-Syn-mEos2 was unaffected by electrical field stimulation at 10 Hz for 99 s in the absence of Dyngo-4a. Following pre-incubation with Dyngo-4a to inhibit endocytosis, the mobility of a α-Syn-mEos2 increased during 10 Hz stimulation to deplete the SV cluster (**Extended Data Fig. 10**). Together, these results demonstrate that the SV cluster is critical for the enrichment of α-Syn at the presynapse.

## Discussion

Using single-particle tracking in motor neurons of live *Drosophila*, we elucidate the mechanism by which cytoplasmic proteins become concentrated at presynapses. Our data support a model in which cytoplasmic proteins are retained within the presynapse through a ‘barrier effect’ at the interface of a dynamic SV scaffold.

We propose that the ‘barrier effect’ emerges from transient and multivalent interactions in a liquid-like network. Proteins inside the network (i.e., the SV scaffold and the cytoplasmic space between SVs) repeatedly form interactions with one another, however, these interactions are transient, allowing components to remain mobile. Near the network’s interface, there is a sharp drop in the density of partners a protein can interact with. To leave the network a protein must break multiple interactions, which carries a free-energy cost, thereby favouring internal retention^11^. This manifests as interfacial reflectivity^33,61^. The enrichment within a liquid-like compartment is predicted to be directly linked to single-molecule dynamics governed by this phenomenon^33,61^. Our work demonstrates that interfacial reflectivity governs presynaptic enrichment *in vivo*.

What are the interactions that mediate this effect? α-Syn’s recruitment to the SV scaffold correlates with the affinity of its N-terminus to curved membranes, implicating protein-lipid interactions with SVs. Since α-Syn remains mobile within the peripheral cluster, any interactions that it has with SVs are likely transient. *In vitro*, the N-terminus of α-Syn rapidly converts between bound and unbound states with small unilamellar vesicles, with bound states estimated to last ∼200 ms^52^. *In vivo*, this conversion may be even faster because of competing interactions, which could explain why protein-lipid interactions do not manifest as an increased immobile fraction of α-Syn molecules at the peripheral cluster. α-Syn’s central NACore region also mediates interactions responsible for the ‘barrier effect’, as deletion of the NACore impaired the recruitment of α-Syn^ΔNAC^ to the peripheral cluster. The hydrophobic NACore region seems to be critical for α-Syn’s ability to undergo homophilic phase separation *in vitro*^37,41,42^. β-Syn, the homologue of α-Syn, principally lacks residues 73-83 that constitutes part of the NACore (residues 68-78) and does not spontaneously phase separate^37,41,42^. Therefore, homophilic protein-protein interactions mediated by the NACore could constitute another interaction. However, we cannot rule out the possibility that the NACore region provides another protein-lipid interaction. Indeed, it has previously been proposed that the N-terminus and NAC domain interact with small unilamellar vesicles independently and the two regions could multivalently bridge two vesicles that are up to 15 nm apart in a ‘double-anchor’ system^62^ (albeit the NAC region binds to small unilamellar vesicles at ∼18 folder lower affinity; N-terminus *K_d_* ∼5.2 µM and NAC region *K_d_* ∼88.9 µM^63^). Regardless, N-terminus and NACore provide multivalent interactions. Moreover, recent work has demonstrated that interactions between α-Syn and VAMP2, mediated by α-Syn’s C-terminus, can mediate phase separation *in vitro*, and induce the association of α-Syn with clusters of vesicles^64^. This has been supported by a separate group, who show that VAMP2/α-Syn/SV-mimics undergo cooperative phase separation *in vitro*, and that deletion of α-Syn’s C-terminus results in a loss of α-Syn from presynapses^65^. Together, α-Syn is capable of multivalent interactions which may underly its experience of a ‘boundary effect’.

A competing interpretation to our proposed model is ‘molecular piggybacking’. Since the diffusion of individual SVs are reflected, an α-Syn molecule tightly associated with an SV would macroscopically appear to ‘bounce’, even in the absence of its own genuine preferential reflectivity at the scaffold interface. However, this interpretation is at odds with several of our results: (1) that α-Syn diffuses almost an order of magnitude faster than SVs, (2) long-lived pausing or slowing that might indicate long-term vesicle association events in the peripheral cluster are not obvious since the fraction of immobile α-Syn molecules are similar in both the peripheral cluster and central axoplasm, and (3) the modest reduction in diffusion coefficient of α-Syn in the peripheral cluster compared to the central axoplasm, cannot, by itself, explain the extent to which α-Syn is enriched in the peripheral cluster. Therefore, despite the necessity of α-Syn’s interaction with SVs for enrichment, this interaction does not result in ‘immobilizing’ events. Rather, this manifests as reflectivity at the interface via short-lived and repeated interactions with SVs, that are not long enough for α-Syn to inherit the low mobility of SVs.

Expectations about the behaviours of single proteins at liquid-liquid interfaces have been predicted^33,61^, but to the best of our knowledge, our work provides the first quantification of the single-molecule dynamics of a protein and a small organelle at an *in vivo* liquid-liquid interface. We anticipate that this principle may apply across other liquid-like structures in the cell, offering a physical mechanism to explain how proteins become locally concentrated while remaining highly mobile and hence functionally available.

## Methods

### Ethical statement

All experimental procedures using mice were conducted under the guidelines of the Australian Code of Practice for the Care and Use of Animals for Scientific purposes and were approved by The University of Queensland (UQ) Animal Ethics Committee (2020/AE000439, 2020/AE000379 and 2023/AE000169). Mice used in this study were C57BL/6 strain wild type. Mice were maintained in a 12-hour light/dark cycle and housed in a PC2 facility with *ad libitum* access to food and water at the Queensland Brain Institute. Both male and female mice were used for all experiments.

### Primary hippocampal neuron cultures

#### Cultures

Primary hippocampal neurons were obtained from C57BL/6 embryonic day-16 mice. ∼45,000 neurons were seeded either onto poly-L-lysine (1 mg/mL poly-L-lysine in borate buffer) coated 29 mm or 35 mm glass-bottom dishes with 10 mm micro-wells (Cellvis, #D29-10-1.5-N or #D35-10-1-N). The neurons were plated in Neurobasal medium (Gibco, #21103-049) supplemented with 5% fetal bovine serum (Hyclone), 2% GlutaMAX (Gibco, #35050061) and 50 U/mL penicillin/streptomycin (Invitrogen, #15140122). The medium was changed to serum-free Neurobasal medium supplemented with 2% B27 (Gibco, #17504-044) 5 to 6 hours post-seeding, and the medium was thereafter supplemented every week.

#### Transduction and transfection

Primary hippocampal neurons were transduced on DIV 3 with an α-Syn KO lentivirus (pLP17 LentiCRISPRV2(LCV2)) at a multiplicity of infection of 5, see ref.^66^. Transduced primary neurons were subsequently transfected with α-Syn-mEos2, and for dual-colour experiments, co-transfected with VAMP2-pHluorin. Transfections were performed on DIV 14 using Lipofectamine® 2000 (Invitrogen, #11668019) as per manufacturer’s instructions.

#### Imaging buffers

During imaging, neurons were incubated in either low K+ resting imaging buffer (145 mM NaCl, 5.6 mM KCl, 2.2 mM CaCl_2_, 0.5 mM MgCl_2_, 5.6 mM D-glucose, 0.5 mM ascorbic acid, 0.1% BSA, 15 mM HEPES, pH 7.4) or phenol free neurobasal media supplemented with 2% GlutaMAX (Gibco, #35050061), 50 U/mL penicillin/streptomycin (Invitrogen, #15140122) and 2% B27 (Gibco, #17504-044).

### Single-molecule microscopy of primary hippocampal neurons

DIV 21 neurons expressing α-Syn-mEos2 (and for dual-color imaging VAMP2-pHluorin) were used for imaging. Single-molecule microscopy was performed using an iLas2 ring-TIRF illumination system (Roper Scientific) mounted on a Nikon Ti-E inverted microscope with a Nikon CFI Apo TIRF 100x oil 1.49 NA objective and Perfect Focus System (Nikon Corporation), as described previously^31^. The system is fitted with two Evolve 512 Delta EMCCD cameras (Teledyne Photometrics) mounted on a TwinCam LS dual-emission image splitter (Cairn Research). For dual-colour imaging, TetraSpeckTM microspheres (Invitrogen-Thermo Fisher Scientific, 0.1 μm) were used for alignment of the transmitted and reflected ports. Timelapse movies (5000 frames) were acquired at 50 Hz using Metamorph (v7.7.8, Molecular Devices, CA, USA).

#### sptPALM

A 405 nm laser (1.5 to 5%; 100 mW Vortran Laser Technology) was used to photoconvert α-Syn-mEos2, and a 561 nm (70%; 150 mW Cobolt Jive) laser was used to simultaneously excite the photoconverted single molecules. To isolate the signal from autofluorescence and background signals, a double-beam splitter (LF488/561-A-000, Semrock, NY, USA) and a double-band emitter (FF01-523/610-25, Semrock, NY, USA) were used.

#### Subdiffractional tracking of internalized molecules (sdTIM)

sdTIM was performed for single-molecule microscopy of recycling vesicles as described previously^7^. Briefly, anti-GFP nanobodies tagged with Atto-647 (Synaptic Systems) were diluted in low K+ imaging buffer at a concentration of 100 pM. Neurons transfected with VAMP2-pHluorin were incubated with the nanobody containing buffer, and were stimulated for 30 s at 10 Hz (100 mA) with the field stimulation insert (described below) to initiate vesicle recycling. The nanobody containing buffer was removed and neurons were washed 2-3 times with low K+ imaging buffer. Cells were incubated for 10 min (37 °C and 5% CO_2_) before being imaged. SptPALM and sdTIM were simultaneously performed for dual-color single-molecule microscopy of α-Syn-mEos2 and VAMP2-pHluorin.

#### 1,6-HD

1,6-HD (1%) (Sigma-Aldrich-Merck, #240117) was prepared in supplemented phenol free neurobasal media. Neurons were incubated with 1,6-HD for 5 min and thereafter imaged in the presence of 1,6-HD for up to 15 minutes.

#### Dyngo-4a and electrical stimulation

Neurons were incubated with 30 µM Dyngo-4a diluted in supplemented phenol free neurobasal medium for 15 min, washed 2-3 times, and imaged immediately after. Neuronal stimulation was performed with 1 ms (100 mA) pulses delivered at 10 Hz for the duration of the acquisition using a field stimulation insert (RC-21BRFS; Warner Instruments, Holliston, MA) and PanLab Channel Digital Stimulator (Warner Apparatus; LE12106 1).

### *Drosophila* stocks and rearing conditions

*Drosophila* stocks were reared on standard yeast-sugar-agar food vials at room temperature (22-25 °C). The parental generation and larval offspring used for experiments were reared in an incubator set at 25 °C prior to microscopy, unless otherwise stated. *Drosophila* stocks used are reported in **Supplementary Table 4**. Generation of *Drosophila* stocks used are reported in **Supplementary Note 2**.

### Microscopy preparation of *Drosophila*

Dissections were performed on crawling third instar larvae bathed in modified HL3 medium (110 mM NaCl, 5 mM KCl, 10 mM MgCl_2_, 10 mM NaHCO_3_, 5 mM trehalose, 30 mM sucrose, 5 mM HEPES, pH 7.2) as described previously^21,67^. A polydimethylsiloxane gel (PDMS; SYLGARD^TM^ 184 Silicone Elastomer Kit) was used as the base for the dissections and microscopy. Briefly, larvae were immobilized on the PDMS gel via inserting a dissection pin between the two posterior spiracles at the tail of the animal, and another between the two anterior spiracles, in front of the mouth hooks. A longitudinal incision was made along the length of the larvae, internal organs were removed, and two dissection pins were placed on each lateral side to pin down the body wall creating a hexagonal silhouette.

### Confocal laser-scanning microscopy of *Drosophila*

Confocal images were acquired using a Plan-Apochromat 63x/1.4 NA oil-immersion objective (Carl Zeiss) on a confocal laser-scanning microscope (LSM 710; Carl Zeiss) built around an Axio Observer Z1 body (Carl Zeiss) and equipped with two internal gallium arsenide phosphide (GaAsP) photomultiplier tubes (PMTs) and three standard PMTs. The system was controlled using Zeiss Zen Black (v2012). 488 nm, 561 nm and 633 nm lasers were used for excitation. PDMS gel with a dissected larva was inverted onto a glass-bottom dish (Cellviz, #D29-20-1.5N) and mounted. Confocal images used for intensity profile analyses were acquired in a *z*-stack with: ×2 scan zoom, 0.07 × 0.07 µm *xy* pixel size, 0.74 µm axial intervals, 1024 × 1024 pixels, 6.27 µs pixel dwell time, 1 airy unit pinhole, unidirectional scanning, ×1 averaging, 16 bit. Confocal images used for boundary contour timelapse were acquired in a timelapse with: ×2 scan zoom, 0.07 × 0.07 µm *xy* pixel size, subset of 1024 × 1024 pixels with a rectangular region of interest (ROI) introduced, 5 s frame time, 6.27 µs pixel dwell time, 1 airy unit pinhole, unidirectional scanning, ×2 averaging, 16 bit.

#### Line-scanning FCS

FCS measurements were acquired on the LSM 710 system described above, using a LCI Plan-Neofluar 63x/1.3 NA water-immersion objective (Carl Zeiss). A 100,000-frame acquisition was made over 120 s across a 4.22 µm region across a bouton with settings: ×32 scan zoom, 0.066 × 0.066 µm *xy* pixel size, 1.2 ms time intervals, 64 pixels across, 4 µs pixel dwell time, unidirectional scanning, ×1 averaging, 16 bit. For auto-correlation analysis custom-written MATLAB scripts were used, as described previously^22^. Briefly, the first 20,000 lines were removed to avoid initial photobleaching, and the remaining line scans were detrended with a moving average. Central and peripheral regions of the bouton were determined manually, and the apparent local diffusion coefficients were determined via fitting the averaged autocorrelation function carpet (within each region) with a one-component diffusion model using a nonlinear least squares method. The point spread function was calibrated using a Rhodamine 110 (20 nM) solution.

### Transmission electron microscopy of *Drosophila*

Third instar larvae were dissected in cold HL3 and fixed for 2 hr at room temperature (4% paraformaldehyde, 1% glutaraldehyde in 0.1 M Na-cacodylate buffer, pH 7.4). Samples were fixed further ON at 4 °C. After washing in 0.1 M Na-cacodylate buffer and afterwards with ddH2O, the larvae were osmicated for 2h (2% OsO4) and stained with 2% uranyl acetate for 1.5 hr. After a final washing step, the larvae were dehydrated with solutions of increasing ethanol concentration (30%, 50%, 70%, 90% and twice with 100%) followed by two 10 min incubation steps with propylene oxide. The larvae then were infiltrated with Agar 100 resin (Laborimpex). Finally, the larvae were embedded in Agar 100 resin and placed in an oven for 48h at +60°C. 70 nm sections were cut on an ultramicrotome (EM UC7, Leica). The sections were imaged using a JEM-1400 transmission electron microscope (Jeol) at 80 keV.

### Single-molecule microscopy of *Drosophila*

Single-particle tracking acquisitions were made using an α-Plan-Apochromat 100×/1.46 NA oil-immersion objective (Carl Zeiss) on an Elyra PS.1 (Carl Zeiss) built around a Axio Observer.Z1 motorized inverted microscope platform (Carl Zeiss) equipped with an iXon 897 EMCCD (Andor Technology), as previously described^20,21^. Acquisitions of up to 15000 frames at 20 ms (45.45 Hz) or 30 ms (31.25 Hz) exposures with ∼2 ms dead time were made using a ×1 scan zoom at 0.1 × 0.1 µm *xy* pixel size over a 256 × 256 pixel region. mEos3.2 was simultaneously illuminated with 405 nm and 561 nm lasers. 405 nm laser power was adapted manually (0.001 to 0.01%) to optimize photoactivation for sparse single-molecule detections. 561 nm laser power remained constant during acquisition (10 to 80%) but was adjusted before acquisitions. For labelling with Syt1-Halo, we incubated larvae with 0.2 nM - 20 nM of JF549 HaloTag-ligand for up to 20 min in modified HL3 solution. For some acquisitions, Syt1-Halo was dual-labelled with JF646 HaloTag-ligand to aid in identification of boutons. Larvae were washed with HL3 prior to imaging to remove excess ligand. JF549 HaloTag was illuminated with 561 nm laser (< 10%). For single-molecule acquisitions, the collimator position and illumination angle were adjusted to maximize signal-to-noise ratio for each acquisition.

### Shibire^ts1^ Experiments

Shi^ts1^ larvae were maintained at 18 °C prior to imaging, males were exclusively used for these experiments. Larvae were dissected at RT, and were treated with high K^+^ solution (at RT or < 37 °C) for 1 min, and washed 3× with modified HL3 solution (at RT or < 37 °C). Following washes, larvae were incubated with 200 nM JF646 Halo-tag ligand at 18 °C or < 37 °C (humidified water bath chamber) for 10 min, washed 3× with modified HL3 solution and imaged at 24/32 °C for 5-30 min.

### Single-particle trajectory analyses

#### Localization and tracking

For α-Syn-mEos3.2, single-molecule detections were localized using wavelet-based segmentation, and trajectories were computed using a simulated annealing-based tracking algorithm with PALMTracer (v2.1.0.28228) on MetaMorph (Molecular Devices). For Syt1-Halo, single-molecule detections were localized using Laplacian of Gaussian detector, and trajectories were computed using Linear Assignment Problem tracker with TrackMate (v7.6.1) on ImageJ (Fiji, NIH; v1.54 and v1.53). For α-Syn-mEos3.2 and Syt1-Halo maximum frame-to-frame linking was 500 nm (∼5 pixels) and 300 nm (∼3 pixels), respectively.

#### Drift correction

Single-molecule microscopy acquisitions with visually obvious drift were drift-corrected using SharpViSu^68^, as described previously^69^. Acquisitions with significant, uncorrectable drift were excluded.

#### ROIs for primary hippocampal neurons

ROIs were hand drawn around synaptic boutons, defined by morphology and pHluorin/mEos2 intensities (trajectories ≥ 8 frames were used for all respective analyses).

#### ROIs for NMJ

ROIs were hand drawn around synaptic boutons and at the peripheral cluster-axoplasm boundary, defined by morphology and single-molecule detection density.

#### Apparent diffusion coefficients

Apparent diffusion coefficients were estimated by fitting MSD-Δt plot (first four points) with: MSD(Δt) = 4*D*Δt + *a* (where *D* is the diffusion coefficient, *a* is the y-intercept, and Δt is the lag time). Apparent diffusion coefficients were calculated for trajectories ≥ 5 detections in length.

#### Immobile fraction

The immobile fraction of molecules was determined using a *D* threshold (*D_min_* = [0.1 µm]^2^ / [4 × 4 × 0.022 s] ≈ 0.028 µm^2^/s), such than an immobile molecule explores an area inferior to the detector’s spatiotemporal resolution, as done before^23^. With this *D_min_*, ∼91% of α-Syn-mEos3.2 molecules were classified as immobile following PFA-fixation (4%) (*D* ≈ 0.01 μm^2^/s, **Extended Data Fig. 2g**).

#### TRamWAy inference

Vector fields of local directional bias were inferred from single-molecule trajectories using TRamWay (v0.6.9)^25^. All trajectories, regardless of detection length, were utilized to infer local directional bias (using *DF* inference) allowing for increased statistical power within local hexagonally tessellated bins. Inference maps computed from boutons without a peripheral cluster lacked local and directed diffusional bias (**Extended Data Fig 2i**).

#### Trajectory angles

We defined a width to the boundary, as the interface appears ‘fuzzy’ when inspected with super-resolved localization maps (since detections are accumulated over time at a fluctuating boundary). We defined the width of the near-boundary region as: maximum particle tracking frame-to-frame distance × 0.5. Custom-written python scripts were used for analysis of trajectory angles near the peripheral cluster-axoplasm boundary:

1. Regions of interest were created by dilating a hand-drawn cluster-axoplasm boundary with shapely.buffer(), applying it once for the *near-boundary* region and a second time for the *distal* region.
2. For every point (p1) in a trajectory, the step angle was calculated, using the preceding point (p0), and following point (p2).
3. For each p0→p1 step, the direction of the trajectory with respect to the interface (*towards* or *away*) was determined by using a combination of shapely.distance() and shapely.contains().
4. Those p1 points that were inside the boundary region, or whose p0→p1 step crossed the central side of the boundary region were used for analysis.

#### p_Exit_

To estimate *p_Exit_*, we filtered by trajectories who started within the *near-boundary* region. From these trajectories, we determined the percentage of trajectories that ended more central (*percC*), more peripheral (perc*P*), or within the *near-boundary* region. *p*_Exit_ was estimated using the formula, *p_Exit_ = (2 × percC) / (percC + percP)*.

#### saSPT

Apparent diffusion coefficients distributions were additionally estimated using saSPT (v0.4.0)^24^, trajectories ≥ 2 detections in length were used^24^. StateArrayDataset was used for batch processing with [pixel_size = 0.1 µm, frame_interval = 0.022 s, diff_coefs = np.logspace(−2, 2, 400)], and other parameters were default.

### Monte Carlo simulation

The volumetric mesh of an axonal segment with presynaptic nerve terminal which formed a synapse onto a dendritic spine was reconstructed using Reconstruct^70^ (v1.1.0.0) from ssTEM (3.8 nm × 3.8 nm × 50 nm resolution) of the hippocampal area CA1^15^ available through the Open NeuroData Registry (https://registry.opendata.aws/open-neurodata). The axonal segment was reconstructed to ∼3.5 µm in length, yielding a model synapse density comparable to that measured in rodent cortical neurons (1 synapse per 3.5 µm axon)^71,72^. The synaptic volume was reconstructed from, and defined as, the volume along the axon occupied by the cluster of SVs. The volumetric meshes were repaired in MeshLab (v2023) and Trimesh (v4.1.7) to remove non-manifold vertices and faces and fill holes. The specified synaptic volume was 0.02 µm^3^ (axis-aligned bounding box of 0.32 µm × 0.32 µm × 0.55 µm), comparable to previously reported values of the synaptic volume occupied by SVs^73^. The volumetric mesh of a representative MN13-Ib bouton was reconstructed using custom-written python scripts from a z-stack acquisition of MN13-Ib bouton in live *Drosophila* expressing Syt1-EGFP (LSM 900; Carl Zeiss; ×6 scan zoom, 0.043 × 0.043 µm xy pixel size, 0.08 µm axial intervals, 397 × 397 pixels, 1.96 µs pixel dwell time, unidirectional scanning, ×4 averaging, 16 bit, Airyscan Mode). The mesh was simplified (decimated) and repaired using MeshLab (v2023). The specified synaptic volume was 6.1 µm^3^ (axis-aligned bounding box of 3.2 µm × 2.3 µm × 3 µm). We specified the axonal volume (11.0 µm^3^) as the vedo.shapes.ConvexHull() of the synaptic volume.

A Monte-Carlo method was used. At simulation start, 1000 particles were placed randomly in the axonal mesh (unless otherwise stated). Particle diffusion was simulated via independent random walks. For each step in a walk, the Gaussian distribution with *σ* = √2*DΔt*, where D is the diffusion coefficient (µm^2^/s) and Δ*t* is the simulation time step (s), was sampled along *x*, *y* and *z* axes. Rejection sampling was used to model the constraint of the axonal boundary.

The simulations contain only 4 base parameters: (1) the reconstructed axonal volume, (2) the reconstructed presynaptic volume, (3) simulation timestep (Δ*t* = 0.0005 s), and (4) number of particles (n_p_ = 1000), all of which remained constant across simulation paradigms. For each simulation paradigm, 2 or 3 additional parameters are introduced:

i. Molecular hindrance: if a particle is inside the synaptic volume, it has diffusion coefficient D_Synapse_, if a particle is outside the synaptic volume, it has diffusion coefficient D_Axon_. D_Synapse_ and D_Axon_ values were varied across a parameter grid of values [0.01, 0.018, 0.032, 0.058, 0.105, 0.190, 0.342, 0.616, 1.110, 2].
ii. Transient immobilizations: a particle has diffusion coefficient D_∀_ regardless of its location, but if a particle is inside the synaptic volume, it has probability p_On_ of becoming immobilized and probability p_Off_ of becoming released from immobilization. D_∀_ was set to 0.616 µm^2^/s, and p_On_ and p_Off_ values were varied across a parameter grid of values [0, 0.001, 0.005, 0.01, 0.025, 0.05, 0.1, 0.25, 0.5, 1].
iii. Confining boundary: a particle has diffusion coefficient D_∀_ regardless of its location, but if a particle is inside the synaptic volume, it has probability p_Exit_ of exiting the synaptic volume if an exiting step is made (if it fails to exit, a non-exiting step is made by rejection sampling). D_∀_ was explored across [0.01, 0.018, 0.032, 0.058, 0.105, 0.190, 0.342, 0.616, 1.110, 2] and p_Exit_ was explored across [0, 0.001, 0.005, 0.01, 0.025, 0.05, 0.1, 0.25, 0.5, 1].

Simulations were run until the spatial distributions of particles reached (or were close to reaching) equilibrium. The number of timesteps (i.e., the total length of simulation in time) are reported in **Supplementary Table 2** and **Supplementary Table 3** and range from 2.5 s to 20 s. Simulation timestep (Δ*t* = 0.0005 s) was chosen to ensure (1) diffusive motion was unaffected by significant degree of rejection sampling by the axonal boundary across the range of diffusion coefficients sampled and (2) efficient simulation runtimes (see **Supplementary Note 1**, **Supplementary Table 1**, and **Supplementary Fig 1**). Number of particles (n_p_ = 1000) was chosen to ensure stable estimates at equilibrium^1^. Fold enrichment was determined as (density in synapse) / (density in axon), where units for density are particles/µm^3^.

### Other analyses

#### Statistics

SciPy (v1.13.1) and Pingouin (v0.5.3) in Python (v3.9.16) were used for statistical analyses. Matplotlib (v3.7.1) and Seaborn (v0.13.2) were used for data visualization. Outlier removal was performed for sptPALM analyses in data obtained from hippocampal neurons. In this case, all data points lying two standard deviations from the mean were considered outliers and removed. No outlier removal was performed for all other statistical analyses.

#### Timelapse contours

Timelapse contours of the axoplasm-cluster boundary were generated using custom-written python scripts. Raw fluorescence images were smoothed to remove noise (2D Gaussian filter, σ = 0.7 to 0.8 pixels) and contours were created with an intensity threshold that decreased over time to account for photobleaching. Contours were visually inspected to confirm accuracy. Contours on the exterior of the bouton remained stable.

#### Intensity line profiles

Regions of interest (ROIs) were generated in ImageJ (Fiji, NIH; *v1.54* and *v1.53*). ROIs were linear lines drawn across the diameter of the bouton; the experimenter was blinded from the experimental condition during ROI creation (except for two comparisons: mEos3.2 vs α-Syn-mEos3.2 and EGFP vs Syt1-EGFP). Raw intensity profiles across the ROI were obtained using ProfilePot() and smoothed to remove noise (1D Gaussian filter, σ = 1 pixel) for proper alignment and interpolated and normalized using custom-written python scripts. Intensity integration was performed between the central region of the intensity profile (−0.1 to 0.1; normalized units of distance, where centre of the aligned line profile is the origin) to calculate central integrated intensity, and the sum of the peripheral regions of the intensity profile (−0.3 to −0.2 and 0.2 to 0.3) to calculate peripheral integrated intensity. A ratio of peripheral to central intensity was generated from these values.

## Acknowledgements

We thank all past and present members of the Meunier and Millard laboratories for valuable discussions. The super-resolution imaging was carried out at the Queensland Brain Institute’s (QBI’s) Advanced Microscopy Facility. We thank Dr. Rumelo Amor, Dr. Andrew Thompson and Dr. Arnaud Gaudin for their technical support for microscopy experiments. We thank Jef Swerts for EM images of *Drosophila* motor neurons. We thank Dr. Tristan Wallis and Helen Y. McGuinness for critical appraisal of the manuscript. We thank Dr. Akihiro Kusumi, Dr. Etienne Herzog, and Dr. Dietmar Oelz for valuable discussions. EGFP-synapsin lines were kindly provided by Dr. Troy Littleton. The VAMP2-pHluorin plasmid was kindly provided by Dr. James Rothman. Dyngo-4a was kindly provided by Dr. Phil Robinson. Computational simulations were supported by resources provided by The University of Queensland Research Computing Centre’s Bunya supercomputer, with funding from The University of Queensland, Brisbane, Australia.

## Contributions

K.K., F.A.M, A.T.B. and S.S.M., conceived the project; K.K., F.A.M., A.T.B., and S.S.M designed the study; K.K performed experiments, analyses, and simulations; S.F.L. performed hippocampal neuron experiments and analyses; K.K., J.S.-V., and E.H. performed line-scanning FCS experiments and analyses; K.K. and A.J.M. performed saSPT analysis; N.S., A.T.B, and P.V. created transgenic fly lines; L.A.P.-R and S.R. prepared α-Syn KO lentivirus; F.A.M. and K.K. acquired funding; K.K. authored the original draft of the manuscript and prepared figures with significant input from F.A.M and S.S.M.; F.A.M., A.T.B and S.S.M. supervised; all authors contributed to writing the manuscript and approved its final form.

