## Supplementary Material for "Single-molecule bouncing at a liquid interface governs presynaptic enrichment"

**Supplementary Note 1. Set up of computational simulation.**

We optimised the parameters used in the three-dimensional synapse simulation based on the following considerations. Rejection sampling was used to model the physical constraint of the axonal plasma membrane; accordingly, the simulation time-step (Δt) should be kept sufficiently small to capture the mobility of the particle moving within the three-dimensional volume of the axon. The standard deviation of the Gaussian distribution ($\sigma = \sqrt{2D\Delta t}$) dictates the degree of the particle's displacement along each of the three dimensions in one simulation step. It is critical that this value, $\sigma$, is small in relation to the size of the axonal mesh to mitigate statistical bias that may arise from over-use of rejection sampling.

To determine which range of values of $\sigma$ result in reliable translational diffusive motion within the reconstructed axon, we plotted the resulting histogram of displacement steps in all three axes for various values of $\sigma$ by adjusting the values of Δt (**Supplementary Fig. 1, Supplementary Table 1**).

Deviation from the normal distribution with $\sigma = \sqrt{2D\Delta t}$ results from the rejection of particle steps at the axon mesh boundary, which becomes more apparent with increased Δt and $\sigma$ (**Supplementary Fig. 1**). At Δt = 0.01 s ($\sigma$ = 0.14 μm), the particle displacements greatly diverge from the expected Gaussian distribution. At Δt = 0.001 s ($\sigma$ = 0.045 μm), the particle displacements slightly diverge from the expected Gaussian distribution. Below Δt = 0.0001 s ($\sigma$ = 0.014 μm) the particle displacements accurately follow the expected Gaussian distribution. There is less deviation from the Gaussian along the z-axis, as this axis is closest to the longitudinal axis of the axon and therefore does not exhibit as much of a degree of confinement. Therefore, for optimal results $\sigma$ should be kept below 0.045 μm. We did so by running simulations at Δt = 0.0005 s, such that our fastest diffusion coefficient simulated ($D$ = 2 μm$^2$/s) resulted in step size sampling from a Gaussian distribution with $\sigma$ = 0.0447 μm. Comparison of apparent diffusion coefficients determined from a two-dimensional projection of three-dimensional simulated trajectories ensured that the simulated model was accurate (**Extended Data Fig. 1**).

To ensure that the simulation is operating as expected, we initialized 1000 particles in the synaptic region and allowed them to diffuse freely (no enrichment model) across different ground-truth diffusion coefficients (**Supplementary Fig. 2**). We measured the apparent diffusion coefficients by fitting the first four points of each trajectory-derived mean square displacement (MSD) curve (using spatial information along the *x* and *z* axes). Apparent diffusion coefficients measured are equivalent to ground truth diffusion coefficients (**Supplementary Fig. 2**).

| $D$ (μm$^2$/s) | Δt (s) | $\sigma$ (μm) |
|---|---|---|
| 1 | 0.01 | 0.141421 |
| 1 | 0.001 | 0.044721 |
| 1 | 0.0001 | 0.014142 |
| 1 | 0.00001 | 0.004472 |
| 1 | 0.000001 | 0.001414 |
| 1 | 0.0000001 | 0.000447 |

**Supplementary Table 1. Values of σ tested in computational simulations.**

The standard deviation of the Gaussian distribution ($\sigma = \sqrt{2D\Delta t}$) for the particle walk steps, was adjusted by changing Δt. *D* = diffusion coefficient, Δt = time-step.

| $D_{Axon}$ (μm²/s) | Steps (#) | Δt (s) |
|---|---|---|
| 0.01 | 40000 | 0.0005 |
| 0.018016 | 36111 | 0.0005 |
| 0.032459 | 32222 | 0.0005 |
| 0.05848 | 28333 | 0.0005 |
| 0.105361 | 24444 | 0.0005 |
| 0.189824 | 20556 | 0.0005 |
| 0.341995 | 16667 | 0.0005 |
| 0.616155 | 12778 | 0.0005 |
| 1.110095 | 8889 | 0.0005 |
| 2 | 5000 | 0.0005 |

**Supplementary Table 2. Number of timesteps used in the computational simulation for each value of $D_{Axon}$ in the computational simulations for the molecular hindrance model.**

Simulations of the molecular hindrance model were run for a specified number of timesteps (at Δt = 0.0005s) that allowed for the particle distribution to approach an equilibrium state.

| D∀ (μm$^2$/s) | Steps (#) | Δt (s) |
|---|---|---|
| 0.01 | 40000 | 0.0005 |
| 0.018016 | 36111 | 0.0005 |
| 0.032459 | 32222 | 0.0005 |
| 0.05848 | 28333 | 0.0005 |
| 0.105361 | 24444 | 0.0005 |
| 0.189824 | 20556 | 0.0005 |
| 0.341995 | 16667 | 0.0005 |
| 0.616155 | 12778 | 0.0005 |
| 1.110095 | 8889 | 0.0005 |
| 2 | 5000 | 0.0005 |

**Supplementary Table 3. Number of timesteps used in the computational simulation for each value of D∀ in the computational simulations for the confining boundary model.**

Simulations of the confining boundary model were run for a specified number of timesteps (at Δt = 0.0005s) that allowed for the particle distribution to approach an equilibrium state.

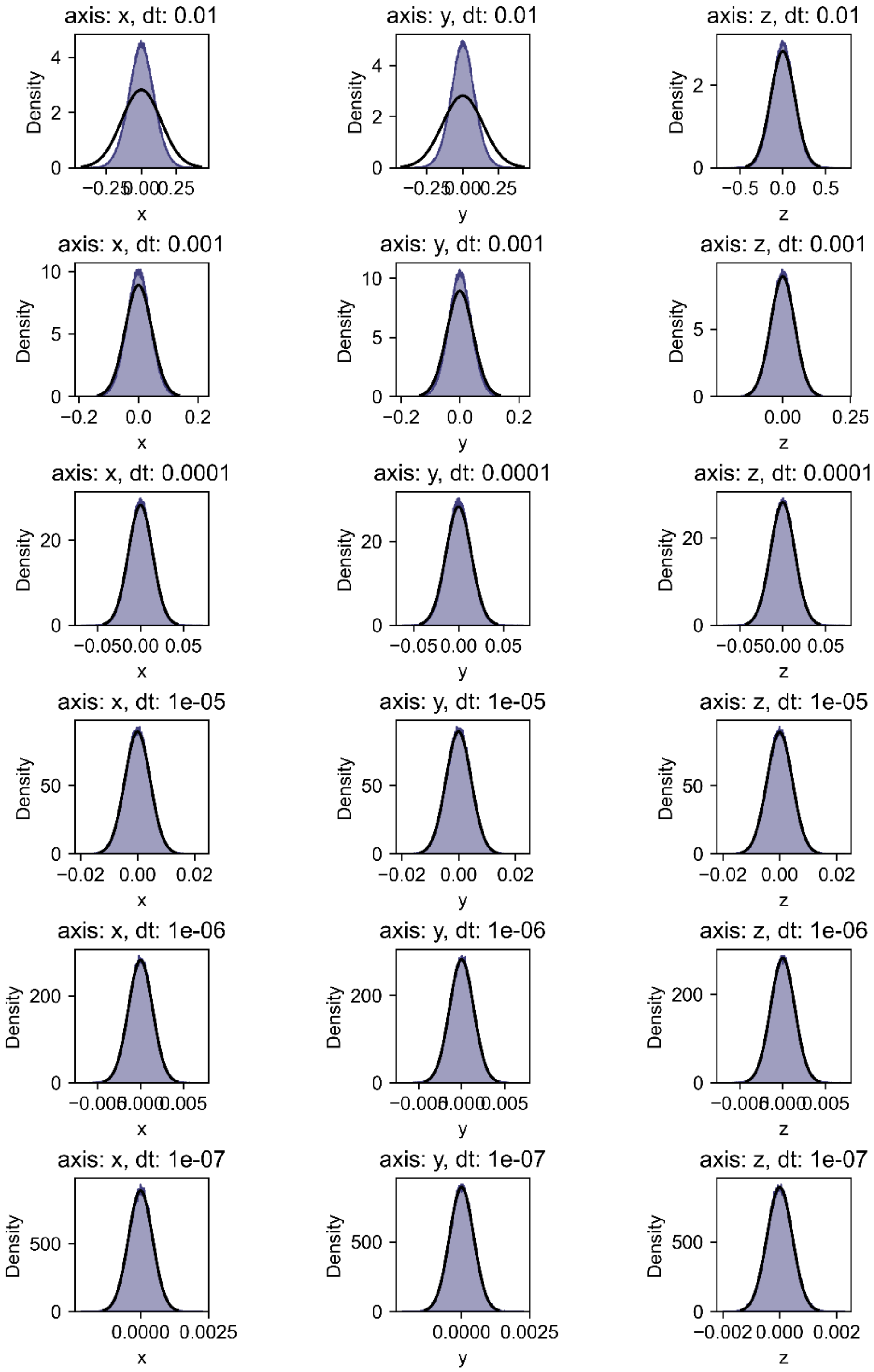


**Supplementary Fig. 1. Smaller simulation step sizes result in more reliable statistics of the translational diffusion of particles.**

Each plot shows the displacement of the simulated particles (blue) and the ground-truth Gaussian distribution ($\sigma = \sqrt{2D\Delta t}$; black) along each of the three-dimensional axes during individual steps (dt = simulation time-step) (**Supplementary Table 1**).

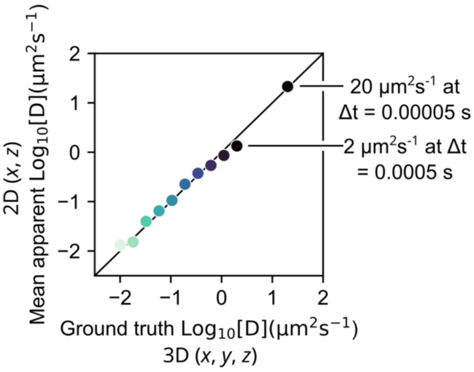


**Supplementary Fig. 2. 3D ground-truth simulated diffusion coefficients are equivalent to those determined by the MSD curve-fitting of simulated trajectory data projected in 2D.**

Simulated 3D trajectory data within the axon (*x*, *y*, *z*) is projected onto 2D (*x*, *z*; where *z* is the long axis), and the apparent single-particle diffusion coefficients are estimated by fitting the first four points of each 2D trajectory-derived mean square displacement (MSD) curve. Each point represents the mean apparent diffusion coefficient from 1000 particles. Identity line is shown in black.

### Supplementary Note 2. Creation of α-Syn *Drosophila* lines

The pUAST-attB plasmid was linearized by restriction digestion with EcoRI and XbaI. The insert was assembled into the linearized backbone using HiFi DNA Assembly (New England Biolabs) with synthesized gBlocks encoding mEos3.2 and SNCA. The SNCA cDNA sequence was codon-optimized for Drosophila melanogaster. A short flexible linker sequence (GGGTGGGS) was introduced between SNCA and mEos3.2 to facilitate proper folding and function of the fusion protein. All constructs were verified by Sanger sequencing (Eurofins genomics). Transgenic flies were generated using the ΦC31 integrase system at the VK00005 attP landing site.

mEos3.2:

GGTGGGGGTACgGGAGGCGGATCCATGAGTGCGATTAAGCCAGACATGAAGATCAAACTCC
GTATGGAAGGCAACGTAAACGGGCACCACTTTGTGATCGACGGAGATGGTACAGGCAAGC
CTTTTGAGGGAAAACAGAGTATGGATCTTGAAGTCAAAGAGGGCGGACCTCTGCCTTTTGCC
TTTGATATCCTGACCACcGCATTCCATTACGGCAACAGGGTATTCGCCAAATATCCAGACAAC
ATACAAGACTATTTTAAGCAGTCGTTTCCTAAGGGGTATTCGTGGGAACGAAGCTTGACTTTCG
AAGACGGGGGCATTTGCAACGCCAGAAACGACATAACAATGGAAGGGGACACTTTCTATAA
TAAAGTTCGATTTTATGGTACCAACTTTCCCGCCAATGGTCCAGTTATGCAGAAGAAGACGCT
GAAATGGGAGCCCTCCACTGAGAAAATGTATGTGCGTGATGGAGTGCTGACGGGTGATATTG
AGATGGCTTTGTTGCTTGAAGGAAATGCCCATTACCGATGTGACTTCAGAACTACTTACAAAG
CTAAGGAGAAGGGTGTCAAGTTACCAGGCGCCCACTTTGTGGACCACTGCATTGAGATTTTA
AGCCATGACAAAGATTACAACAAGGTTAAGCTGTATGAGCATGCTGTTGCTCATTCTGGATTG
CCTGACAATGCCAGACGATAAGGGTACCTCTAGAGGATCTTT

SNCA WT:

AACTCTGAATAGGGAATTGGGcAAacATGGACGTCTTTATGAAGGGACTGAGTAAAGCGAAGG
AGGGTGTGGTGGCGGCCGCAGAAAAAACGAAGCAGGGCGTGGCAGAGGCGGCGGGAA
AGACGAAGGAAGGAGTGCTGTATGTTGGCTCGAAAACCAAGGAAGGCGTCGTGCACGGCG
TTGCCACAGTTGCGGAGAAGACGAAGGAGCAGGTAACCAACGTAGGAGGAGCTGTAGTAA
CAGGCGTAACCGCCGTTGCCCAAAAGACCGTCGAGGGAGCTGGTTCGATAGCCGCTGCT
ACCGGTTTTGTAAAAAAAGATCAGTTGGGGAAAAACGAAGAGGGTGCTCCGCAGGAGGGAA
TCCTGGAAGACATGCCGGTGGACCCAGATAATGAGGCATACGAGATGCCATCCGAAGAAG
GCTACCAGGATTATGAACCAGAGGCAGGTGGGGGTACgGGAGGCGGA

SNCA A30P:

AACTCTGAATAGGGAATTGGGcAAacATGGACGTCTTTATGAAGGGACTGAGTAAAGCGAAGG
AGGGTGTGGTGGCGGCCGCAGAAAAAACGAAGCAGGGCGTGGCAGAGGCGcCGGGAAA
GACGAAGGAAGGAGTGCTGTATGTTGGCTCGAAAACCAAGGAAGGCGTCGTGCACGGCGT
TGCCACAGTTGCGGAGAAGACGAAGGAGCAGGTAACCAACGTAGGAGGAGCTGTAGTAAC
AGGCGTAACCGCCGTTGCCCAAAAGACCGTCGAGGGAGCTGGTTCGATAGCCGCTGCTAC
CGGTTTTGTAAAAAAAGATCAGTTGGGGAAAAACGAAGAGGGTGCTCCGCAGGAGGGAATC

CTGGAAGACATGCCGGTGGACCCAGATAATGAGGCATACGAGATGCCATCCGAAGAAGGC
TACCAGGATTATGAACCAGAGGCAGGTGGGGGTACgGGAGGCGGA

SNCA A53T:

AACTCTGAATAGGGAATTGGGcAAacATGGACGTCTTTATGAAGGGACTGAGTAAAGCGAAGG
AGGGTGTGGTGGCGGCCGCAGAAAAAACGAAGCAGGGCGTGGCAGAGGCGGCGGGAA
AGACGAAGGAAGGAGTGCTGTATGTTGGCTCGAAAACCAAGGAAGGCGTCGTGCACGGCG
TTaCCACAGTTGCGGAGAAGACGAAGGAGCAGGTAACCAACGTAGGAGGAGCTGTAGTAAC
AGGCGTAACCGCCGTTGCCCAAAAGACCGTCGAGGGAGCTGGTTCGATAGCCGCTGCTAC
CGGTTTTGTAAAAAAAGATCAGTTGGGGAAAAACGAAGAGGGTGCTCCGCAGGAGGGAATC
CTGGAAGACATGCCGGTGGACCCAGATAATGAGGCATACGAGATGCCATCCGAAGAAGGC
TACCAGGATTATGAACCAGAGGCAGGTGGGGGTACgGGAGGCGGA

SNCA H50Q:

AACTCTGAATAGGGAATTGGGcAAacATGGACGTCTTTATGAAGGGACTGAGTAAAGCGAAGG
AGGGTGTGGTGGCGGCCGCAGAAAAAACGAAGCAGGGCGTGGCAGAGGCGGCGGGAA
AGACGAAGGAAGGAGTGCTGTATGTTGGCTCGAAAACCAAGGAAGGCGTCGTGCAgGGCG
TTGCCACAGTTGCGGAGAAGACGAAGGAGCAGGTAACCAACGTAGGAGGAGCTGTAGTAA
CAGGCGTAACCGCCGTTGCCCAAAAGACCGTCGAGGGAGCTGGTTCGATAGCCGCTGCT
ACCGGTTTTGTAAAAAAAGATCAGTTGGGGAAAAACGAAGAGGGTGCTCCGCAGGAGGGAA
TCCTGGAAGACATGCCGGTGGACCCAGATAATGAGGCATACGAGATGCCATCCGAAGAAG
GCTACCAGGATTATGAACCAGAGGCAGGTGGGGGTACgGGAGGCGGA

SNCA G51D:

AACTCTGAATAGGGAATTGGGcAAacATGGACGTCTTTATGAAGGGACTGAGTAAAGCGAAGG
AGGGTGTGGTGGCGGCCGCAGAAAAAACGAAGCAGGGCGTGGCAGAGGCGGCGGGAA
AGACGAAGGAAGGAGTGCTGTATGTTGGCTCGAAAACCAAGGAAGGCGTCGTGCACGaCG
TTGCCACAGTTGCGGAGAAGACGAAGGAGCAGGTAACCAACGTAGGAGGAGCTGTAGTAA
CAGGCGTAACCGCCGTTGCCCAAAAGACCGTCGAGGGAGCTGGTTCGATAGCCGCTGCT
ACCGGTTTTGTAAAAAAAGATCAGTTGGGGAAAAACGAAGAGGGTGCTCCGCAGGAGGGAA
TCCTGGAAGACATGCCGGTGGACCCAGATAATGAGGCATACGAGATGCCATCCGAAGAAG
GCTACCAGGATTATGAACCAGAGGCAGGTGGGGGTACgGGAGGCGGA

SNCA ΔNAC:

AACTCTGAATAGGGAATTGGGcAAacATGGACGTCTTTATGAAGGGACTGAGTAAAGCGAAGG
AGGGTGTGGTGGCGGCCGCAGAAAAAACGAAGCAGGGCGTGGCAGAGGCGGCGGGAA
AGACGAAGGAAGGAGTGCTGTATGTTGGCTCGAAAACCAAGGAAGGCGTCGTGCACGGCG
TTGCCACAGTTGCGGAGAAGACGAAGGAGCAGGTAACCAACGTAGGACAAAAGACCGTCG
AGGGAGCTGGTTCGATAGCCGCTGCTACCGGTTTTGTAAAAAAAGATCAGTTGGGGAAAAAC
GAAGAGGGTGCTCCGCAGGAGGGAATCCTGGAAGACATGCCGGTGGACCCAGATAATGAG
GCATACGAGATGCCATCCGAAGAAGGCTACCAGGATTATGAACCAGAGGCAGGTGGGGGT
ACgGGAGGCGGA

| *Type* | *Short name* | *Full genotype* | *Stock #* |
|---|---|---|---|
| driver | OK6-GAL4 | P{w[+mW.hs]=GawB}OK6 | 64199 |
| driver | nSyb-GAL4 | y[1] w[*]; P{w[+m*]=nSyb-GAL4.S}3 | 51635 |
| transgene | UAS-mEos3.2 | UAS-mEos3.2 in VK00005 | this paper |
| transgene | UAS-α-Syn-mEos3.2 | UAS-α-Syn-mEos3.2 in VK00005 | this paper |
| transgene | UAS-α-SynA30P-mEos3.2 | UAS-α-SynA30P-mEos3.2 in VK00005 | this paper |
| transgene | UAS-α-SynG51D-mEos3.2 | UAS-α-SynG51D-mEos3.2 in VK00005 | this paper |
| transgene | UAS-α-SynE46K-mEos3.2 | UAS-α-SynE46K-mEos3.2 in VK00005 | this paper |
| transgene | UAS-α-SynH50Q-mEos3.2 | UAS-α-SynH50Q-mEos3.2 in VK00005 | this paper |
| transgene | UAS-α-SynΔNAC-mEos3.2 | UAS-α-SynΔNAC-mEos3.2 in VK00005 | this paper |
| transgene | UAS-Syt1-Halo | w[*]; P{y[+t7.7] w[+mC]=UAS-Syt1::Halo7}attP40 | 67622 |
| transgene | UAS-Syt1-EGFP | w[*]; P{w[+mC]=UAS-syt.eGFP}2 | 6925 |
| transgene | UAS-EGFP | w[1118]; P{w[+mC]=UAS-EGFP}5a.2 | 5431 |
| transgene | UAS-EGFP-Synapsin | (3rd Chromosome) | Vasin et al., (2014) |
| mutant | Shits1 | w[1118] shi[1] | 7068 |
| RNAi | y[1] v[1]; P{y[+t7.7] v[+t1.8]=TRiP.JF01829}attP2 | | 25811 |
| RNAi | y[1] v[1]; P{y[+t7.7] v[+t1.8]=TRiP.JF01918}attP2 | | 25879 |
| RNAi | y[1] v[1]; P{y[+t7.7] v[+t1.8]=TRiP.JF02049}attP2 | | 26024 |
| RNAi | y[1] v[1]; P{y[+t7.7] v[+t1.8]=TRiP.JF02590}attP2 | | 27279 |
| RNAi | y[1] v[1]; P{y[+t7.7] v[+t1.8]=TRiP.JF02613}attP2 | | 27304 |
| RNAi | y[1] v[1]; P{y[+t7.7] v[+t1.8]=TRiP.JF02634}attP2 | | 27484 |
| RNAi | y[1] v[1]; P{y[+t7.7] v[+t1.8]=TRiP.JF02652}attP2 | | 27502 |
| RNAi | y[1] v[1]; P{y[+t7.7] v[+t1.8]=TRiP.JF02667}attP2 | | 27517 |
| RNAi | y[1] v[1]; P{y[+t7.7] v[+t1.8]=TRiP.JF02729}attP2 | | 27652 |
| RNAi | y[1] v[1]; P{y[+t7.7] v[+t1.8]=TRiP.JF02793}attP2 | | 27713 |
| RNAi | y[1] v[1]; P{y[+t7.7] v[+t1.8]=TRiP.JF02841}attP2/TM3, Sb[1] | | 28007 |
| RNAi | y[1] v[1]; P{y[+t7.7] v[+t1.8]=TRiP.JF02901}attP2 | | 28064 |
| RNAi | y[1] v[1]; P{y[+t7.7] v[+t1.8]=TRiP.JF02994}attP2 | | 28358 |
| RNAi | y[1] v[1]; P{y[+t7.7] v[+t1.8]=TRiP.JF03130}attP2/TM3, Sb[1] | | 28510 |
| RNAi | y[1] v[1]; P{y[+t7.7] v[+t1.8]=TRiP.JF03139}attP2 | | 28712 |
| RNAi | y[1] v[1]; P{y[+t7.7] v[+t1.8]=TRiP.JF02471}attP2 | | 29312 |
| RNAi | y[1] v[1]; P{y[+t7.7] v[+t1.8]=TRiP.JF03336}attP2 | | 29401 |
| RNAi | y[1] v[1]; P{y[+t7.7] v[+t1.8]=TRiP.JF03417}attP2 | | 31983 |
| RNAi | y[1] sc[*] v[1] sev[21]; P{y[+t7.7] v[+t1.8]=TRiP.HMS00403}attP2 | | 32408 |
| RNAi | y[1] sc[*] v[1] sev[21]; P{y[+t7.7] v[+t1.8]=TRiP.HMS00928}attP2 | | 33971 |
| RNAi | y[1] sc[*] v[1] sev[21]; P{y[+t7.7] v[+t1.8]=TRiP.HMS01131}attP2 | | 34655 |
| RNAi | y[1] sc[*] v[1] sev[21]; P{y[+t7.7] v[+t1.8]=TRiP.HMS00248}attP2 | | 34944 |
| RNAi | y[1] sc[*] v[1] sev[21]; P{y[+t7.7] v[+t1.8]=TRiP.HMS00459}attP2 | | 34958 |
| RNAi | y[1] sc[*] v[1] sev[21]; P{y[+t7.7] v[+t1.8]=TRiP.HMS01414}attP2 | | 35004 |
| RNAi | y[1] sc[*] v[1] sev[21]; P{y[+t7.7] v[+t1.8]=VALIUM20-mCherry.RNAi}attP2 | | 35785 |
| RNAi | y[1] sc[*] v[1] sev[21]; P{y[+t7.7] v[+t1.8]=TRiP.GL00249}attP2 | | 36061 |
| RNAi | y[1] sc[*] v[1] sev[21]; P{y[+t7.7] v[+t1.8]=TRiP.HMS01724}attP40 | | 38274 |
| RNAi | y[1] sc[*] v[1] sev[21]; P{y[+t7.7] v[+t1.8]=TRiP.HMS01736}attP40 | | 38283 |
| RNAi | y[1] v[1]; P{y[+t7.7] v[+t1.8]=TRiP.HMS01932}attP40 | | 39014 |
| RNAi | y[1] v[1]; P{y[+t7.7] v[+t1.8]=TRiP.HMS01937}attP40 | | 39019 |
| RNAi | y[1] v[1]; P{y[+t7.7] v[+t1.8]=TRiP.HMS01939}attP40 | | 39021 |
| RNAi | y[1] sc[*] v[1] sev[21]; P{y[+t7.7] v[+t1.8]=TRiP.HMS01993}attP40/CyO | | 39073 |
| RNAi | y[1] sc[*] v[1] sev[21]; P{y[+t7.7] v[+t1.8]=TRiP.HMS02011}attP2 | | 40845 |
| RNAi | y[1] v[1]; P{y[+t7.7] v[+t1.8]=TRiP.HMS02042}attP40 | | 40875 |
| RNAi | y[1] sc[*] v[1] sev[21]; P{y[+t7.7] v[+t1.8]=TRiP.HMS02442}attP40 | | 42017 |
| RNAi | y[1] sc[*] v[1] sev[21]; P{y[+t7.7] v[+t1.8]=TRiP.HMS02636}attP40 | | 42943 |
| RNAi | y[1] sc[*] v[1] sev[21]; P{y[+t7.7] v[+t1.8]=TRiP.HMS02728}attP40 | | 44014 |
| RNAi | y[1] v[1]; P{y[+t7.7] v[+t1.8]=TRiP.HMC02936}attP40 | | 44541 |
| RNAi | y[1] sc[*] v[1] sev[21]; P{y[+t7.7] v[+t1.8]=TRiP.HMS02846}attP2 | | 44551 |
| RNAi | y[1] v[1]; P{y[+t7.7] v[+t1.8]=TRiP.HMJ03119}attP40 | | 50901 |
| RNAi | y[1] v[1]; P{y[+t7.7] v[+t1.8]=TRiP.HMJ21316}attP40 | | 53951 |
| RNAi | y[1] v[1]; P{y[+t7.7] v[+t1.8]=TRiP.HMC02373}attP2 | | 55241 |
| RNAi | y[1] sc[*] v[1] sev[21]; P{y[+t7.7] v[+t1.8]=TRiP.HMC04412}attP40 | | 56972 |
| RNAi | y[1] v[1]; P{y[+t7.7] v[+t1.8]=TRiP.HMS04480}attP40 | | 57046 |
| RNAi | y[1] sc[*] v[1] sev[21]; P{y[+t7.7] v[+t1.8]=TRiP.HMS04495}attP40 | | 57298 |
| RNAi | y[1] sc[*] v[1] sev[21]; P{y[+t7.7] v[+t1.8]=TRiP.HMC04936}attP40 | | 57743 |
| RNAi | y[1] v[1]; P{y[+t7.7] v[+t1.8]=TRiP.HMJ21977}attP40 | | 58085 |
| RNAi | y[1] sc[*] v[1] sev[21]; P{y[+t7.7] v[+t1.8]=TRiP.HMC06275}attP2 | | 66338 |
| RNAi | y[1] sc[*] v[1] sev[21]; P{y[+t7.7] v[+t1.8]=TRiP.HMS05447}attP40 | | 66981 |
| RNAi | y[1] sc[*] v[1] sev[21]; P{y[+t7.7] v[+t1.8]=TRiP.HMS06092}attP40 | | 82983 |

**Supplementary Table 4. *Drosophila* lines used**

Stock # as indicated in Bloomington *Drosophila* Stock Center.

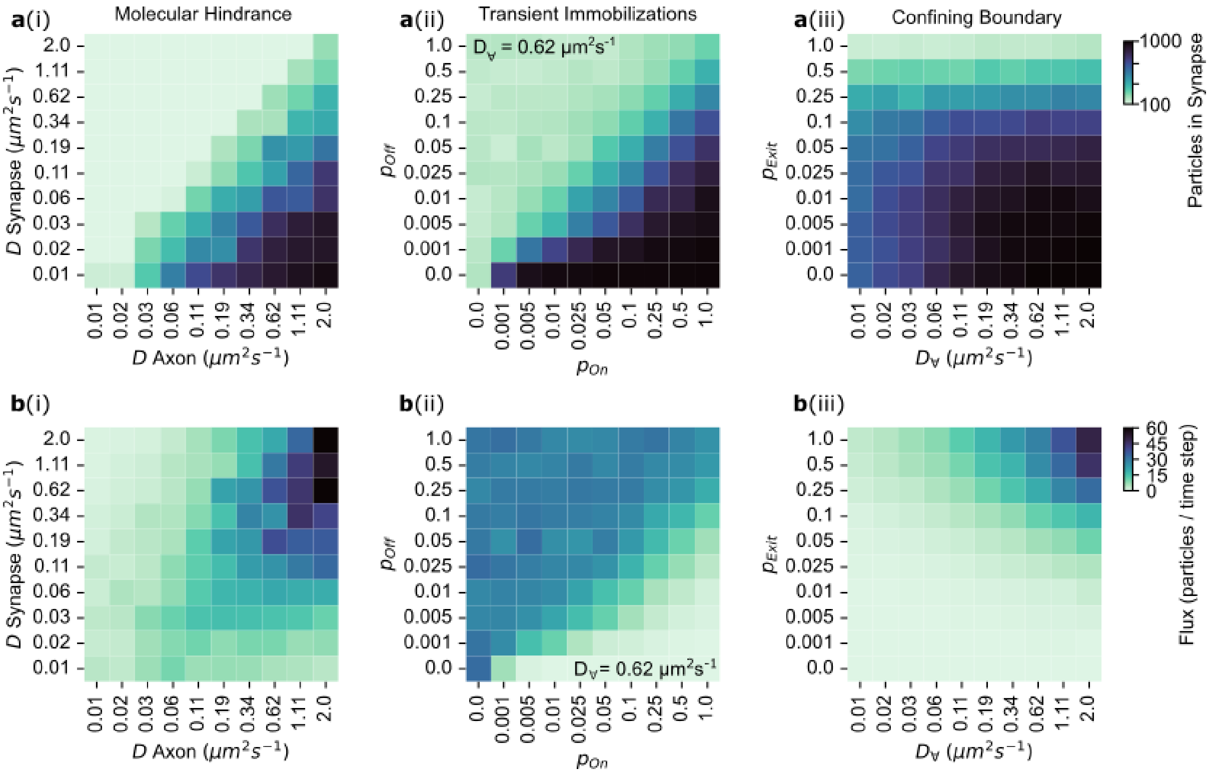


**Extended Data Fig. 1 Presynaptic particle enrichment and flux across matrix of input parameters for models of enrichment.**

(**a**, **b**) Each model of enrichment was run with a matrix of input parameters (**Methods, Supplementary Note 1, Supplementary Table 2, Supplementary Table 3**). (**a**) Matrix heatmaps show the number of particles located within the synaptic volume at (or approaching) simulation equilibrium, for the given input parameters of each model. (**b**) Heatmaps show particle flux across the synaptic volume (particles entering synaptic volume per timestep) at, or close to, simulation equilibrium. (i) Molecular hindrance, (ii) transient immobilizations and (iii) confining boundary models.

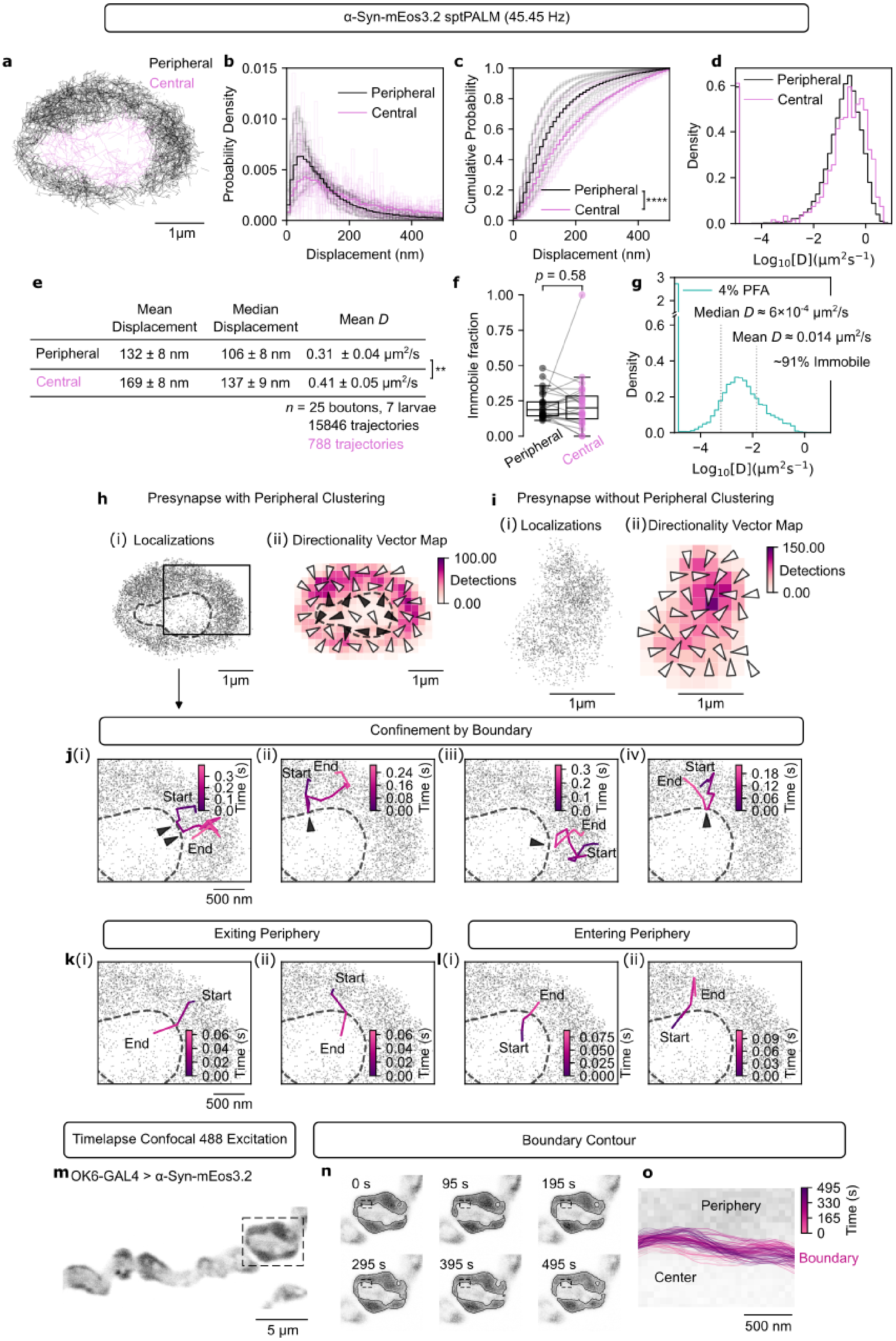

α-Syn-mEos3.2 sptPALM (45.45 Hz)
a
Peripheral
Central
1μm
b
Probability Density
Displacement (nm)
Peripheral
Central
c
Cumulative Probability
Displacement (nm)
Peripheral
Central
****
d
Density
Peripheral
Central
Log10[D](μm2s−1)
e
Mean Displacement
Median Displacement
Mean D
Peripheral 132 ± 8 nm 106 ± 8 nm 0.31 ± 0.04 μm2/s
Central 169 ± 8 nm 137 ± 9 nm 0.41 ± 0.05 μm2/s
**
n = 25 boutons, 7 larvae
15846 trajectories
788 trajectories
f
p = 0.58
Immobile fraction
Peripheral
Central
g
4% PFA
Median D ≈ 6×10-4 μm2/s
Mean D ≈ 0.014 μm2/s
~91% Immobile
Density
Log10[D](μm2s−1)
h
Presynapse with Peripheral Clustering
(i) Localizations
(ii) Directionality Vector Map
100.00
Detections
0.00
1μm
i
Presynapse without Peripheral Clustering
(i) Localizations
(ii) Directionality Vector Map
150.00
Detections
0.00
1μm
Confinement by Boundary
j(i)
(ii)
(iii)
(iv)
Start
End
Time (s)
500 nm
Exiting Periphery
Entering Periphery
k(i)
(ii)
l(i)
(ii)
Start
End
Time (s)
500 nm
Timelapse Confocal 488 Excitation
Boundary Contour
m OK6-GAL4 > α-Syn-mEos3.2
5 μm
n
0 s
95 s
195 s
295 s
395 s
495 s
o
Periphery
Boundary
Center
Time (s)
500 nm

**Extended Data Fig. 2. Mobility of α-Syn-mEos3.2 within intra-synaptic regions of MN13-Ib boutons.**

(**a**) Representative sptPALM trajectories of α-Syn-mEos3.2 in a MN13-Ib bouton; steps made within the central axoplasm are shown in pink. (**b**) Probability distribution and (**c**) cumulative probability distribution of α-Syn-mEos3.2 trajectory step lengths within the central axoplasm and peripheral cluster (two-sided Kolmogorov-Smirnov test). (**d**) Distribution of apparent diffusion coefficients (*D*) of α-Syn-mEos3.2 trajectories in the central axoplasm (pink) or peripheral cluster (black). (**e**) Table of mean displacement, median displacement, and mean diffusion coefficient within the central axoplasm and the peripheral cluster (two-sided paired Wilcoxon signed-rank test on *D*). (**f**) Immobile fraction of molecules ($n$ = 25 boutons from 7 larvae; two-sided paired Wilcoxon signed-rank test). (**g**) Distribution of *D* in MN13-Ib from larva fixed with 4% PFA, indicating that ~91% become immobile after fixation ($n$ = 4 NMJs, from 1 larva). (**h**, **i**) sptPALM localization map (i), and inference map overlayed with detection heatmap (ii) of MN13-Ib bouton in **a**, which showed peripheral clustering (**h**), and representative bouton with no apparent peripheral clustering (**i**); dashed line represents boundary between central axoplasm and peripheral cluster for **h**; vectors near the boundary of the central axoplasm and the peripheral cluster in **h** are coloured black. (**j**) Representative trajectories of α-Syn-mEos3.2 with hindered motion (black arrows) at the boundary between the central axoplasm and the peripheral cluster (inset from **h**). (**k**, **l**) Representative trajectories of α-Syn-mEos3.2 crossing the boundary between the central axoplasm and peripheral cluster; (**k**) exiting, and (**l**) entering the peripheral cluster. (**m**) Representative confocal image of MN13-Ib expressing α-Syn-mEos3.2 used for time-lapse imaging. (**n**) Bouton indicated in **m** acquired over a 500 s acquisition; contours are shown around peripheral high-intensity region. (**o**) Contours coloured in time from region indicated in **n**, the interface between the peripheral cluster and central axoplasm appears to undulate over time. Data shown as mean ± SEM; $p$-values indicated as: ****, $p \leq 0.0001$; **, $p \leq 0.01$.

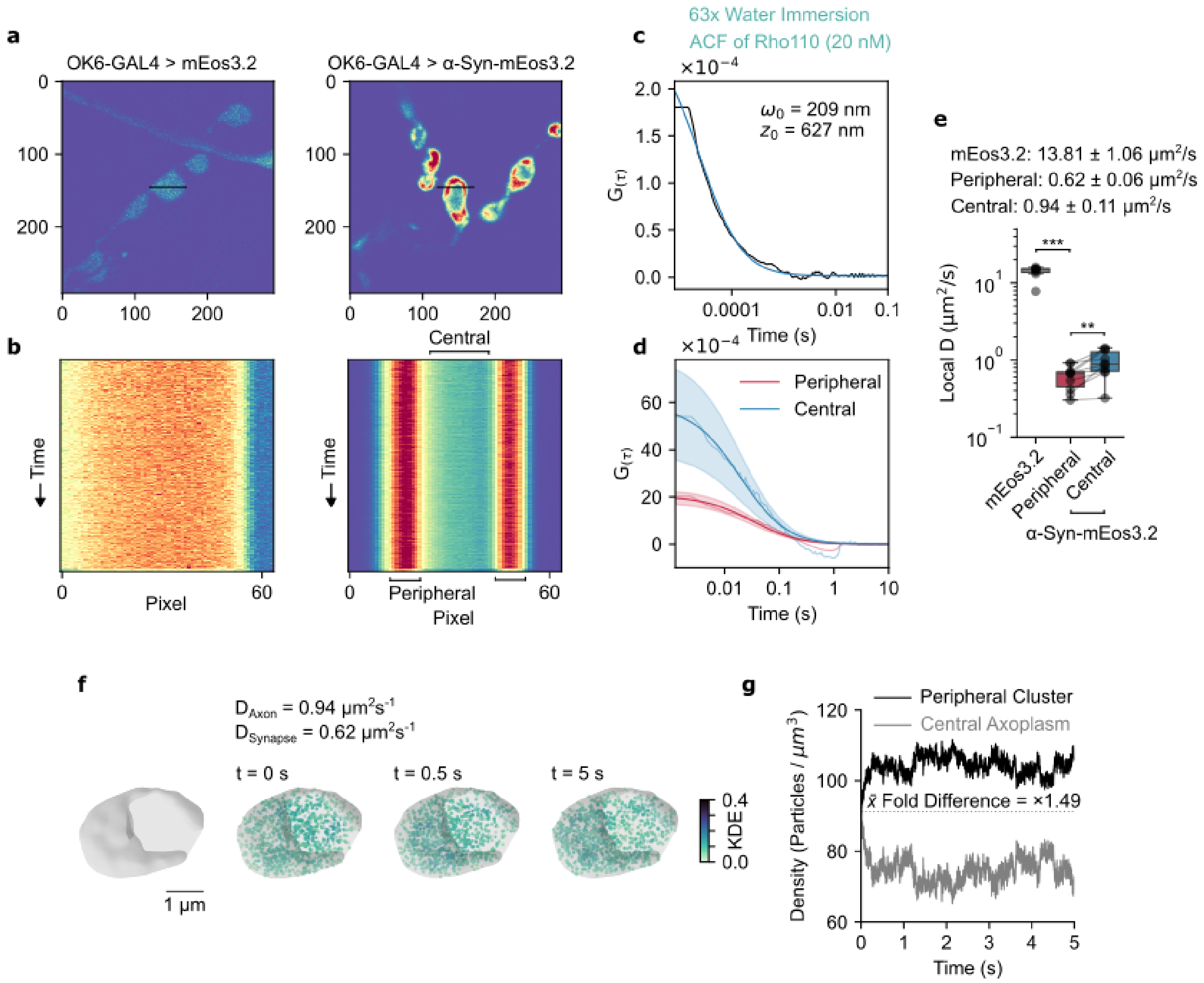


**Extended Data Fig. 3. Line-scanning FCS within MN13-Ib boutons.**

(**a**) Representative confocal images of MN13-Ib expressing (left) mEos3.2 and (right) α-Syn-mEos3.2 (xy-axis in pixels, 92 nm/pixel), with (black) the selected line across a bouton used for line-scanning FCS measurements. (**b**) For line-scanning FCS measurements, a line-scan acquisition (64 × 1 pixels; 66 nm/pixel) was made repeatedly over 120 s at 1.2 ms intervals. The fluctuations in the intensity within each pixel over time results from fluorescent proteins diffusing in and out of the laser beam waist each time the line is scanned. The fluctuations in intensity are represented as a carpet (x-axis is pixel position, and y-axis is time) for (left) mEos3.2 and (right) α-Syn-mEos3.2. (**c**) Calibration measurement for determining point spread function dimensions, autocorrelation function for Rhodamine 110 fitted to a one-component model. (**d**) The autocorrelation function for α-Syn-mEos3.2 derived from correlating intensity fluctuations within each pixel with itself for all possible delay times (τ), grouped by those pixels within the central region, and peripheral region as represented in **b** (central, $n$ = 10 regions; peripheral, $n$ = 20 regions; from 3 larvae). (**e**) Estimated local diffusion coefficients from autocorrelation function (mEos3.2, $n$ = 7 boutons from 2 larvae; α-Syn-mEos3.2, $n$ = 10 boutons from 3 larvae; peripheral-central, two-sided paired Wilcoxon signed-rank test; mEos3.2-peripheral, two-sided Mann-Whitney U test). (**f**) Simulation within 3D-reconstructed MN13-Ib bouton, with different diffusion coefficients within peripheral ($D_{Synapse}$) and central ($D_{Axon}$) regions as specified. (**g**) Density of particles within peripheral and central regions over time from **f**, simulations result in the fold enrichment of only ~1.49. Data shown as mean ± SEM; *p*-values indicated as: ***, $p \leq 0.001$; **, $p \leq 0.01$.

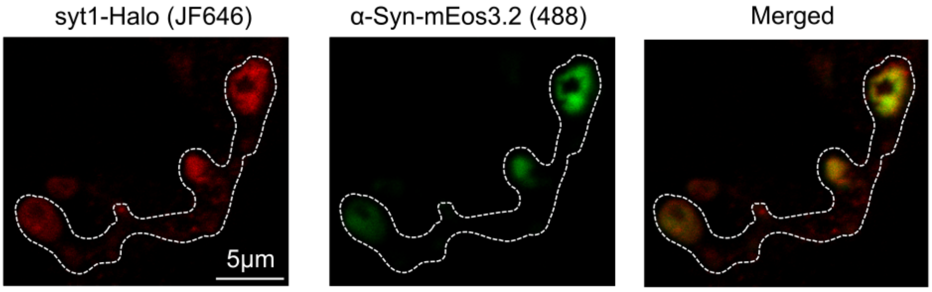


**Extended Data Fig.4. α-Syn colocalizes with the peripheral SV cluster at the MN13-Ib bouton.**

Representative confocal images of MN13-Ib from live *Drosophila* larva expressing Syt1-Halo (JF646; 200 nM) and α-Syn-mEos3.2.

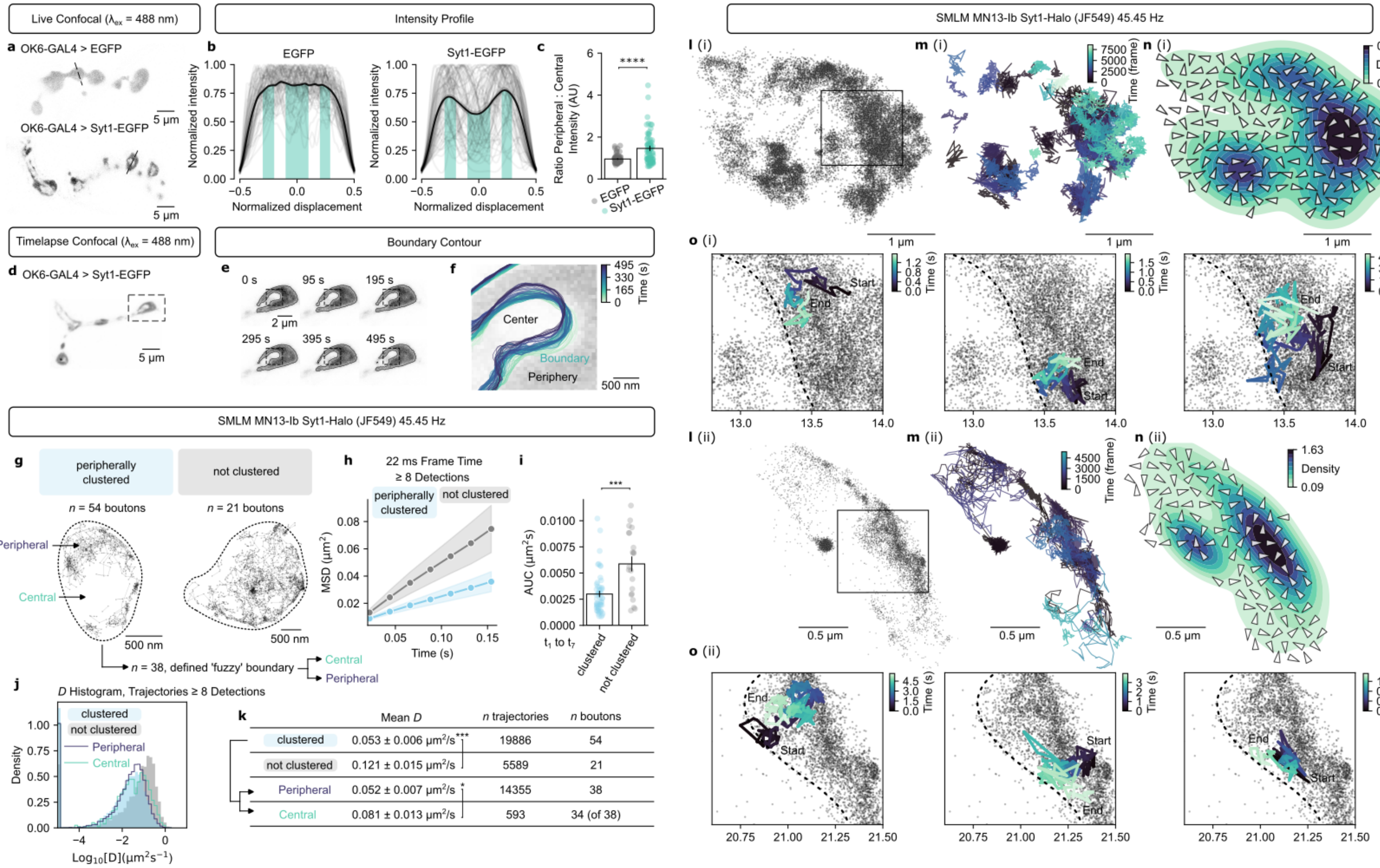

Live Confocal (λex = 488 nm)
a OK6-GAL4 > EGFP
OK6-GAL4 > Syt1-EGFP
5 µm
Intensity Profile
b EGFP
Syt1-EGFP
Normalized intensity
Normalized displacement
c Ratio Peripheral : Central Intensity (AU)
EGFP
Syt1-EGFP
****
Timelapse Confocal (λex = 488 nm)
d OK6-GAL4 > Syt1-EGFP
Boundary Contour
e 0 s 95 s 195 s 295 s 395 s 495 s
2 µm
f Center
Boundary
Periphery
500 nm
Time (s)
SMLM MN13-Ib Syt1-Halo (JF549) 45.45 Hz
g peripherally clustered
not clustered
n = 54 boutons
n = 21 boutons
Peripheral
Central
500 nm
n = 38, defined 'fuzzy' boundary
Central
Peripheral
h 22 ms Frame Time
≥ 8 Detections
MSD (µm²)
Time (s)
i AUC (µm²s)
t1 to t7
clustered
not clustered
***
j D Histogram, Trajectories ≥ 8 Detections
clustered
not clustered
Peripheral
Central
Density
Log10[D](µm²s⁻¹)
k Mean D | n trajectories | n boutons
clustered 0.053 ± 0.006 µm²/s*** 19886 54
not clustered 0.121 ± 0.015 µm²/s 5589 21
Peripheral 0.052 ± 0.007 µm²/s* 14355 38
Central 0.081 ± 0.013 µm²/s 593 34 (of 38)
SMLM MN13-Ib Syt1-Halo (JF549) 45.45 Hz
l (i)
m (i)
n (i)
Time (frame)
Density
1 µm
o (i)
Start
End
Time (s)
l (ii)
m (ii)
n (ii)
0.5 µm
o (ii)

**Extended Data Fig. 5. SV dynamics in *Drosophila* boutons**

(**a**) Representative confocal images of MN13-Ib from live larvae expressing EGFP (top) and Syt1-EGFP (bottom). (**b**) Profiles of the fluorescence intensity across MN13-Ib boutons expressing EGFP (left) and Syt1-EGFP (right). Intensity profiles across individual boutons (grey) and mean intensity profile (black). (**c**) Ratio of peripheral to central intensity acquired from intensity profiles, by integrating the profiles within the shaded central region and peripheral regions indicated in **b** (GFP, $n = 79$ boutons from 3 larvae; Syt1-EGFP, $n = 82$ boutons from 2 larvae; two-sided Mann-Whitney U test). (**d**) Representative confocal image of MN13-Ib expressing Syt1-EGFP used for time-lapse imaging. (**e**) Presynaptic bouton indicated in **d** acquired over a 500 s acquisition; contours are shown around peripheral high-intensity region. (**f**) Contours coloured in time from the region indicated in **e**. (**g**) Representative Syt1-Halo (JF549) trajectories from boutons (45.45 Hz; OK6-GAL4 > Syt1-Halo). Boutons were manually classified as either peripherally clustered (left) or not clustered (right) based on detection and trajectory distribution. Of the 54 peripherally clustered boutons, 38 had a clearly definable boundary. (**h**) MSD-$\Delta t$ plot. (**i**) Corresponding area under the curve (AUC) of the MSD-$\Delta t$ plot (peripherally clustered, $n = 54$ boutons; not clustered, $n = 21$ boutons; taken from 9 larvae; two-sided Mann-Whitney U test). (**j**) Distribution of MSD-fitting-derived diffusion coefficients ($D$) of Syt1-Halo trajectories from peripherally/not clustered boutons, and peripheral/central region of peripherally clustered boutons ($n$ trajectories, see **k**). (**k**) Mean diffusion coefficients ($n$ boutons, see **k**; peripherally/not clustered, two-sided Mann-Whitney U test; peripheral/central, two-sided paired Wilcoxon signed-rank test). (**l**) Representative single-molecule localization map of Syt1-Halo (JF549) in boutons (OK6-GAL4 > Syt1-Halo at 45.45 Hz; from two independent larvae [i] and [ii]). (**m**) Single-molecule trajectories generated after tracking of localizations in **l** (trajectories with length ≥ 30 detections are shown). (**n**) Vector field of local directional bias inferred from single-molecule trajectories in **m** overlayed on top of the kernel density estimate of single-molecule detections. (**o**) Representative single-molecule trajectories of Syt1-Halo displaying hindered motion at the boundary between the axoplasmic matrix and the peripheral Syt1-Halo cluster from inset shown in **l** (axis values in µm). Data shown as mean ± SEM; $p$-values indicated as: ****, $p \leq 0.0001$; ***, $p \leq 0.001$; *, $p \leq 0.05$.

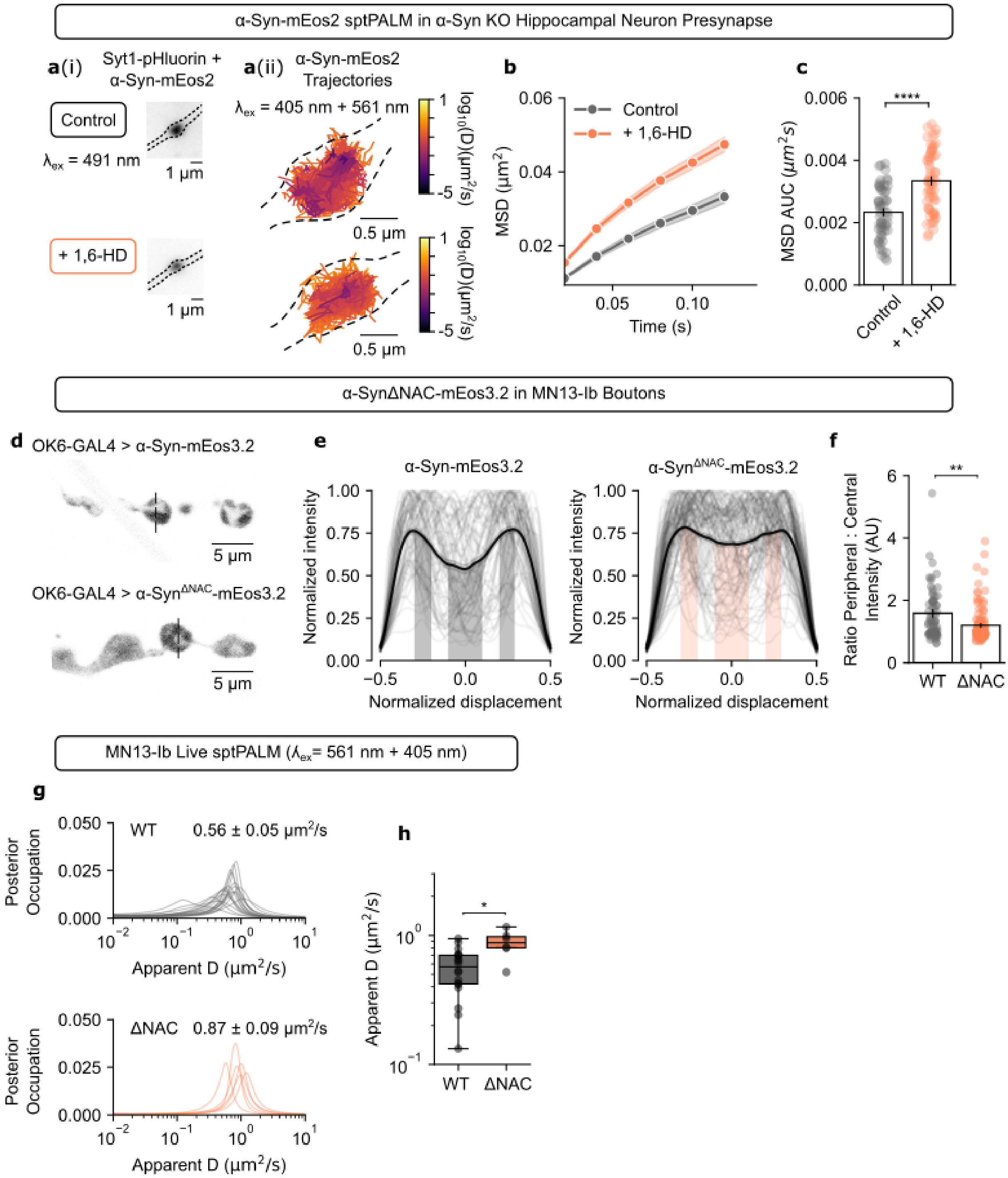
α-Syn-mEos2 sptPALM in α-Syn KO Hippocampal Neuron Presynapse
a(i)
Syt1-pHluorin + α-Syn-mEos2
Control
λex = 491 nm
1 μm
+ 1,6-HD
1 μm
a(ii)
α-Syn-mEos2 Trajectories
λex = 405 nm + 561 nm
0.5 μm
log10(D)(μm²/s)
b
Control
+ 1,6-HD
MSD (μm²)
Time (s)
c
MSD AUC (μm²s)
****
Control
+ 1,6-HD
α-SynΔNAC-mEos3.2 in MN13-Ib Boutons
d
OK6-GAL4 > α-Syn-mEos3.2
5 μm
OK6-GAL4 > α-SynΔNAC-mEos3.2
e
α-Syn-mEos3.2
α-SynΔNAC-mEos3.2
Normalized intensity
Normalized displacement
f
Ratio Peripheral : Central Intensity (AU)
**
WT
ΔNAC
MN13-Ib Live sptPALM (λex= 561 nm + 405 nm)
g
Posterior Occupation
WT 0.56 ± 0.05 μm²/s
ΔNAC 0.87 ± 0.09 μm²/s
Apparent D (μm²/s)
h
Apparent D (μm²/s)
*
WT
ΔNAC

**Extended Data Fig. 6. 1,6-HD disrupts the clustering of α-Syn-mEos2 in cultured hippocampal neuron presynapses and α-Syn$^{\Delta NAC}$-mEos3.2 displays disrupted presynaptic clustering at MN13-Ib boutons.**

(**a**) Syt1-pHluorin and α-Syn-mEos2 at the presynapse of α-Syn KO cultured hippocampal neurons under vehicle control and 1% 1,6-HD treatment. (**a**i) Representative diffraction-limited image of Syt1-pHluorin and α-Syn-mEos2 at the presynapse. (**a**ii) Representative single-molecule trajectories of α-Syn-mEos2 at the presynapse in **a**(i), coloured by diffusion coefficient (top, vehicle control; bottom = 1% 1,6-HD treatment). (**b**) MSD-Δt plot for α-Syn-mEos2 trajectories (of length ≥ 8 detections) in presynaptic boutons in the control and 1% 1,6-HD treated neurons. (**c**) Corresponding area under the curve (AUC) of the MSD-Δt plot for α-Syn-mEos2, each point represents the mean AUC from an individual bouton (Control, *n* = 57 boutons from 2 independent cultures; 1,6-HD, *n* = 60 boutons from 2 independent cultures). (**d**) Representative confocal images of MN13-Ib from live larvae expressing α-Syn-mEos3.2 (top) and α-Syn$^{\Delta NAC}$-mEos3.2 (bottom). (**e**) Profiles of fluorescence intensity across MN13-Ib boutons expressing α-Syn-mEos3.2 (left) and α-Syn$^{\Delta NAC}$-mEos3.2 (right). Intensity profiles of individual boutons (grey) and mean of the profile (black). (**f**) Ratio of peripheral to central intensity computed from intensity profiles in **e** (α-Syn-mEos3.2, *n* = 73 boutons from 3 larvae; α-Syn$^{\Delta NAC}$-mEos3.2, *n* = 117 boutons from 3 larvae; two-sided Mann-Whitney U Test). (**g**) Distribution of diffusion coefficient inferred using saSPT (**Methods**) from single-molecule trajectories of (top) α-Syn-mEos3.2 and (bottom) α-Syn$^{\Delta NAC}$-mEos3.2 (α-Syn-mEos3.2, *n* = 18 NMJs from 8 larvae; α-Syn$^{\Delta NAC}$-mEos3.2, *n* = 6 NMJs from 3 larvae). (**h**) Median apparent diffusion coefficients obtained from saSPT (α-Syn-mEos3.2, *n* = 18 NMJs from 8 larvae; α-Syn$^{\Delta NAC}$-mEos3.2, *n* = 6 NMJs from 3 larvae; two-sided Mann-Whitney U Test). Data shown as mean ± SEM; *p*-values indicated as: ****, $p \leq 0.0001$; **, $p \leq 0.01$; *, $p \leq 0.05$.

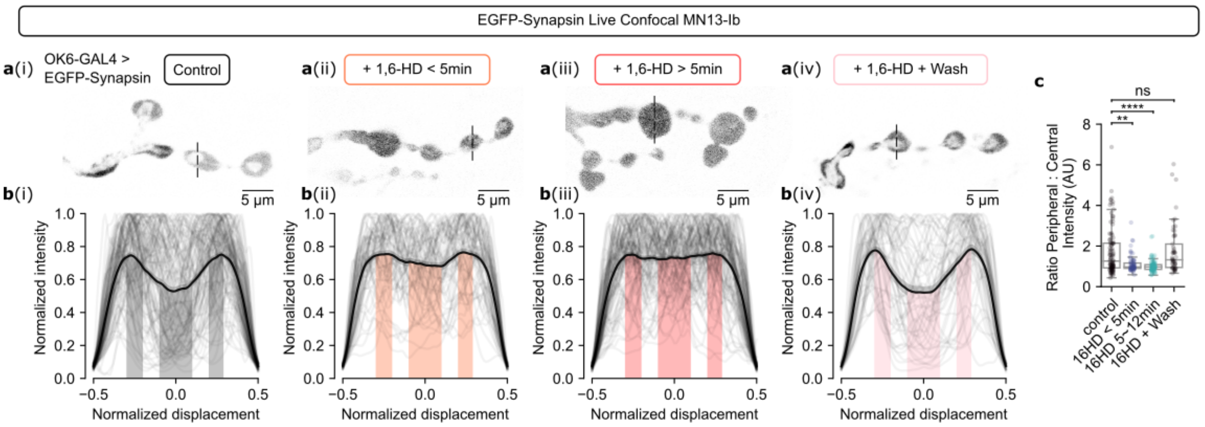


**Extended Data Fig. 7. 4% 1,6-HD abolishes presynaptic clustering of EGFP-Synapsin.**

(**a**) Representative confocal images of MN13-Ib from live *Drosophila* larvae expressing EGFP-Synapsin (OK6-GAL4 > EGFP-Synapsin), under (**a**i) vehicle (modified HL3) control, (**a**ii) addition of 4% 1,6-HD for < 5 min, and (**a**iii) addition of 4% 1,6-HD > 5min (and up to 12 min), and (**a**iv) addition of 4% 1,6-HD for 5 min, followed by modified HL3 wash-off (≥10 min). (**b**) Profiles of fluorescence intensity of EGFP-Synapsin across MN13-Ib boutons, under (**b**i) vehicle (modified HL3) control, (**b**ii) addition of 4% 1,6-HD for < 5 min, and (**b**iii) addition of 4% 1,6-HD > 5min (and up to 12 min), and (**b**iv) addition of 4% 1,6-HD for 5 min, followed by modified HL3 wash-off (≥10 min). Intensity profiles of individual boutons (grey) and mean of the intensity line scans (black). (**c**) Ratio of peripheral/central intensity acquired from intensity profiles (Control, *n* = 113 boutons from 5 larvae; 1,6-HD, *n* = 70 boutons from 7 larvae; 1,6-HD > 5 min, *n* = 110 boutons from 6 larvae; Wash, *n* = 53 boutons from 3 larvae). Data shown as conventional box-and-whisker plots; *p*-values indicated as: **, p ≤ 0.01; ****, *p* ≤ 0.0001; *ns*, not significant *p* > 0.05.

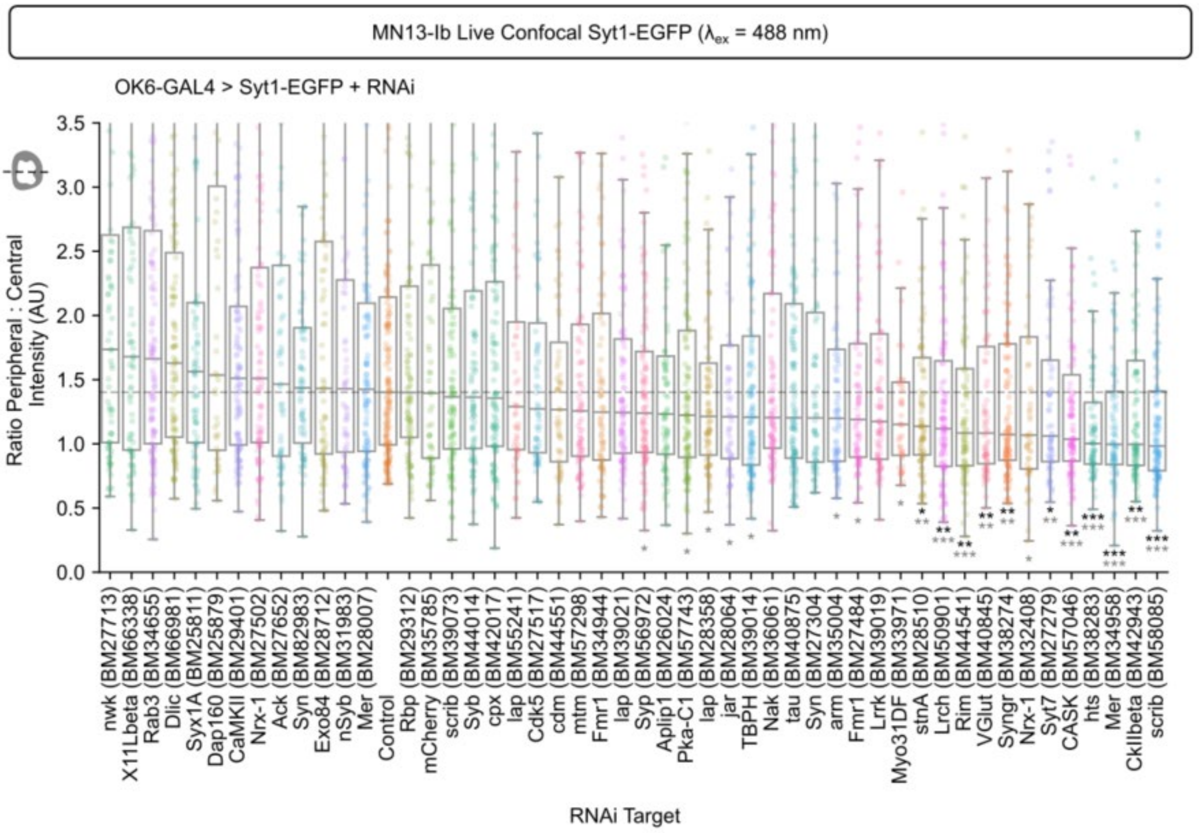


**Extended Data Fig. 8 RNAi-mediated knockdown screen identifies molecular determinants of SV cluster.**

Confocal images of Syt1-EGFP at MN13-Ib boutons were acquired in live *Drosophila* larvae expressing both Syt1-EGFP and dsRNA targeting the candidate gene. Ratio of peripheral to central intensity obtained from Syt1-EGFP intensity profiles across MN13-Ib boutons were obtained for each the dsRNA target. Horizontal dotted line is drawn at the median value of the Control (nSyb-Gal4 > Syt1-EGFP). As a negative control, we used larvae expressing a dsRNA hairpin against mCherry (OK6-GAL4 > Syt1-EGFP + mCherry.RNAi), which exhibited peripheral localisation. Data shown as conventional box-and-whisker plots; *p*-values indicated as: *, $p \leq 0.05$ **, $p \leq 0.01$; ***, $p \leq 0.001$; non-significant $p > 0.05$ comparisons are displayed with no asterisk. Grey* represent uncorrected p-values from two-sided Mann-Whitney U Test, Black* represent p-values from two-sided Mann-Whitney U Test corrected after Benjamini-Hochberg False Discovery Rate correction. Note, two UAS-TRiP lines were performed for: Fmr1, Nrx-1, Syn, lap, scrib, and Mer. The lack of correlation between these RNAi lines may be a result of different dsRNA expression levels.

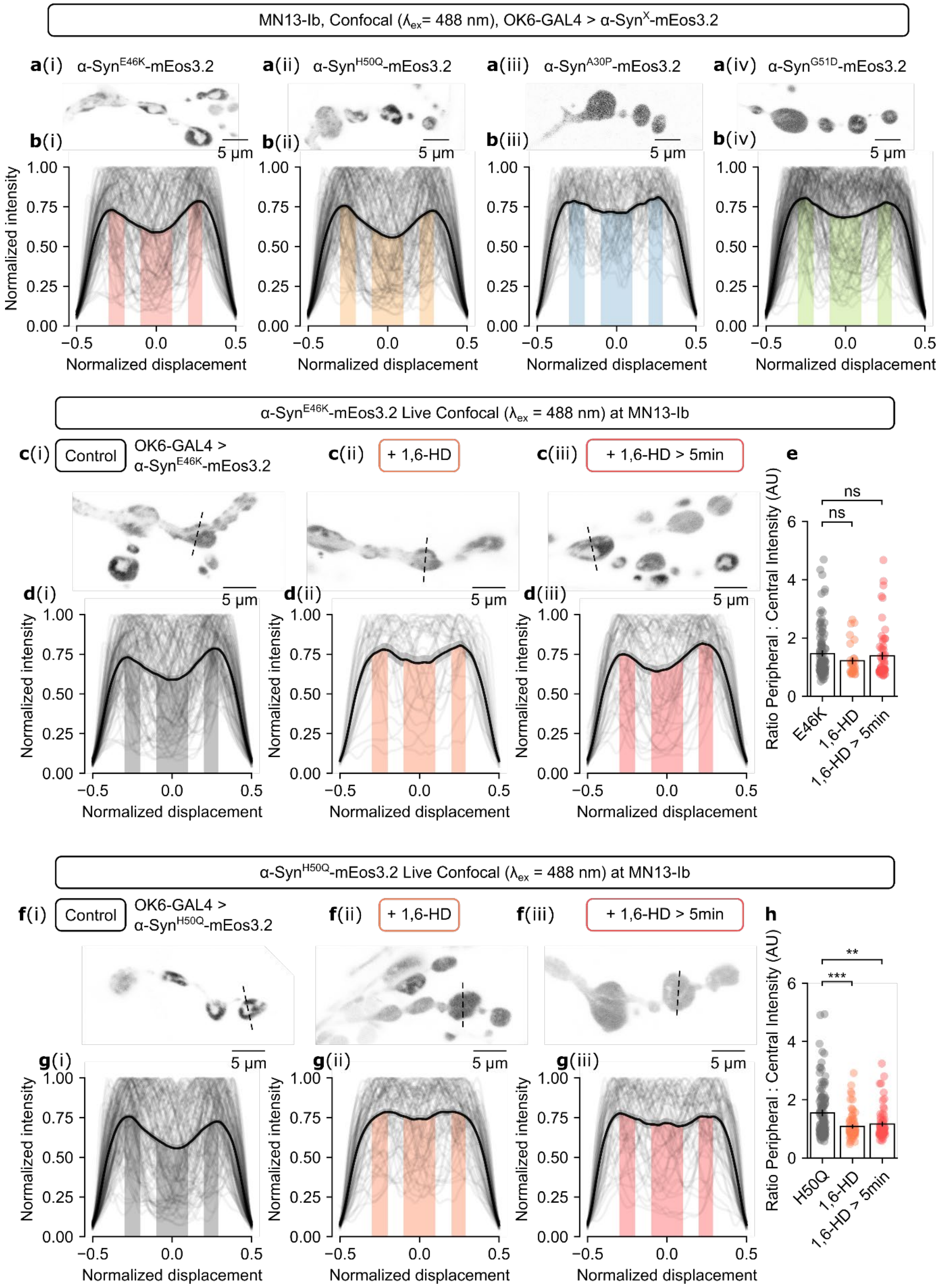
MN13-Ib, Confocal (λex= 488 nm), OK6-GAL4 > α-SynX-mEos3.2
a(i) α-SynE46K-mEos3.2
a(ii) α-SynH50Q-mEos3.2
a(iii) α-SynA30P-mEos3.2
a(iv) α-SynG51D-mEos3.2
b(i) b(ii) b(iii) b(iv)
5 μm
Normalized intensity
Normalized displacement
α-SynE46K-mEos3.2 Live Confocal (λex = 488 nm) at MN13-Ib
c(i) Control OK6-GAL4 > α-SynE46K-mEos3.2
c(ii) + 1,6-HD
c(iii) + 1,6-HD > 5min
d(i) d(ii) d(iii)
e
Ratio Peripheral : Central Intensity (AU)
ns
E46K
1,6-HD
1,6-HD > 5min
α-SynH50Q-mEos3.2 Live Confocal (λex = 488 nm) at MN13-Ib
f(i) Control OK6-GAL4 > α-SynH50Q-mEos3.2
f(ii) + 1,6-HD
f(iii) + 1,6-HD > 5min
g(i) g(ii) g(iii)
h
***
**
H50Q

**Extended Data Fig. 9. MN13-Ib boutons expressing mEos3.2-tagged α-Syn with indicated mutations.**

(**a**) Representative confocal images of MN13-Ib boutons from live $3^{rd}$ instar *D. melanogaster* larvae expressing (**a**i) α-Syn$^{E46K}$-mEos3.2, (**a**ii) α-Syn$^{H50Q}$-mEos3.2, (**a**iii) α-Syn$^{A30P}$-mEos3.2 and (**a**iv) α-Syn$^{G51D}$-mEos3.2 under OK6-GAL4 ($\lambda_{ex}$ = 488 nm). (**b**) Line scan profiles of the fluorescence intensity of (**b**i) α-Syn$^{E46K}$-mEos3.2, (**b**ii) α-Syn$^{H50Q}$-mEos3.2, (**b**iii) α-Syn$^{A30P}$-mEos3.2 and (**b**iv) α-Syn$^{G51D}$-mEos3.2 across MN13-Ib boutons ($\lambda_{ex}$ = 488). Intensity line scans of individual boutons (grey) and mean of the intensity line scans (black). (α-Syn$^{E46K}$-mEos3.2, *n* = 110 boutons from 4 larvae; α-Syn$^{H50Q}$-mEos3.2, *n* = 128 boutons from 5 larvae; α-Syn$^{A30P}$-mEos3.2, *n* = 65 boutons from 3 larvae; α-Syn$^{G51D}$-mEos3.2, *n* = 156 boutons from 6 larvae). (**c, d**) Representative confocal images (**c**) and fluorescence intensity profiles (**d**) of α-Syn$^{E46K}$-mEos3.2 at MN13-Ib boutons under (i) vehicle (modified HL3) control, addition of 4% 1,6-HD for (ii) < 5 min, and (iii) > 5 min. (**e**) Quantification of intensity profiles from **d** (E46K, *n* = 110 boutons from 4 larvae; 1,6-HD, *n* = 33 boutons from 4 larvae, 1,6-HD > 5 min, *n* = 62 boutons from 3 larvae; note E46K data from **Fig. 5**). (**f**, **g**) Representative confocal images (**f**) and fluorescence intensity profiles (**g**) of α-Syn$^{H50Q}$-mEos3.2 at MN13-Ib boutons under (i) vehicle (modified HL3) control, addition of 4% 1,6-HD for (ii) < 5 min, and (iii) > 5 min. (**h**) Quantification of intensity profiles from **g** (H50Q, *n* = 128 boutons from 5 larvae; 1,6-HD, *n* = 79 boutons from 6 larvae, 1,6-HD > 5 min, *n* = 66 boutons from 6 larvae; note H50Q data from **Fig. 5**).

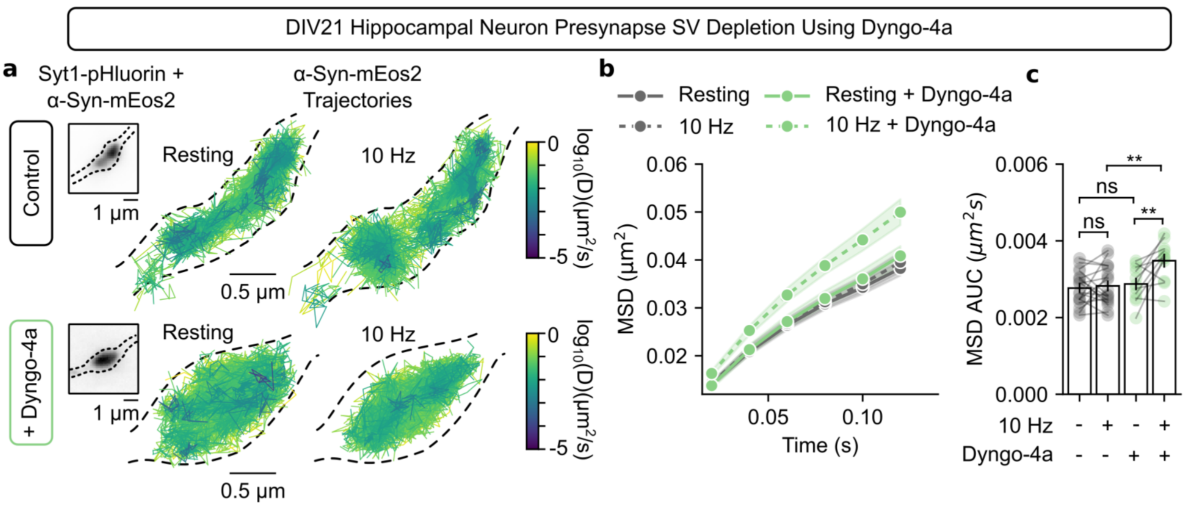


**Extended Data Fig. 10 Effect of Dyngo-4a on α-Syn-mEos2 mobility in primary hippocampal neurons.**

(**a-c**) sptPALM of α-Syn-mEos2 at the cultured hippocampal neuron presynapse (DIV21) with and without Dyngo-4a (30 µM) at 50 Hz acquisition. (**a**) Representative (left) diffraction-limited acquisition of Syt1-pHluorin and α-Syn-mEos2 ($\lambda_{ex}$ = 491 nm) localized within the presynapse and (right) sptPALM single-molecule trajectory map of α-Syn-mEos2 within respective presynapse at resting state and 10 Hz electrical stimulation; experiments were performed (top) without and (bottom) with Dyngo-4a (coloured by diffusion coefficient). (**b**) MSD-Δ*t* plot for α-Syn-mEos2 trajectories localized at the presynapse. (**c**) Corresponding area under the curve (AUC) of the MSD-Δ*t* plot for α-Syn-mEos2, each point represents the mean AUC from synapses in a field of view (Resting and 10 Hz, *n* = 18 acquisitions from 2 biologically independent hippocampal cultures; Resting + Dyngo-4a and 10 Hz + Dyngo-4a, *n* = 11 acquisitions from 3 biologically independent hippocampal cultures). *p*-values indicated as: **; *ns*, not significant $p > 0.05$.
